# A viral toolkit for recording transcription factor-DNA interactions in live mouse tissues

**DOI:** 10.1101/538504

**Authors:** Alexander J. Cammack, Arnav Moudgil, Tomas Lagunas, Jiayang Chen, Michael J. Vasek, Mark Shabsovich, Katherine McCullough, June He, Xuhua Chen, Misha Hooda, Michael N. Wilkinson, Timothy M. Miller, Robi D. Mitra, Joseph D. Dougherty

**Affiliations:** Washington University School of Medicine, Department of Neurology, St. Louis, MO; Washington University School of Medicine, Edison Family Center for Genome Sciences and Systems Biology, St. Louis, MO; Washington University School of Medicine, Department of Genetics, St. Louis, MO; Washington University School of Medicine, Medical Scientist Training Program, St. Louis, MO; Washington University School of Medicine, Department of Psychiatry, St. Louis, MO

## Abstract

Transcription factors (TFs) enact precise regulation of gene expression through site-specific, genome-wide binding. Common methods for TF occupancy profiling, such as chromatin immunoprecipitation, are limited by requirement of TF-specific antibodies and provide only endpoint snapshots of TF binding. Alternatively, TF-tagging techniques, in which a TF is fused to a DNA-modifying enzyme that marks TF binding events across the genome as they occur, do not require TF-specific antibodies and offer the potential for unique applications, such as recording of TF occupancy over time and cell type-specificity through conditional expression of the TF-enzyme fusion. Here we create a viral toolkit for one such method, calling cards, and demonstrate that these reagents can be delivered to the live mouse brain and used to report TF occupancy. Further, we establish a Cre-dependent calling cards system, termed Flip-Excision (FLEX) calling cards and, in proof-of-principle experiments, show utility in defining cell type-specific TF profiles and recording and integrating TF binding events across time. This versatile approach will enable unique studies of TF-mediated gene regulation in live animal models.

## Introduction

Proper cellular development and function is a complex process established by elaborate gene expression networks. These networks are fundamentally regulated by transcription factors (TF), which bind to regulatory elements (RE) across the genome and facilitate gene expression through a variety of mechanisms, including recruitment of transcriptional co-factors and modulation of chromatin state^1^. Extensive efforts to profile TF genome occupancy and identify active REs across the genome have highlighted the profound diversity of TF binding, providing important insights into TF-mediated gene regulation^2–5^. Further, a large portion of genetic variation associated with improper cellular function or disease has been shown to exist in TF-bound REs^3,6–10^, demonstrating the criticality of proper TF binding in maintaining cellular homeostasis.

Several methods exist for profiling TF occupancy across the genome. Antibody-based techniques, such as chromatin immunoprecipitation followed by sequencing (ChIP-seq), and more recently Cleavage Under Targets and Release Using Nuclease (CUT&RUN)^11^ or Tagmentation (CUT&Tag)^12^, are widely used and have provided numerous insights into the cellular functions of TFs^2–4,7^. Notably however, these methods require the availability and individual optimization of TF-specific antibodies, limiting the throughput and breadth of genome-wide TF profiling. Further, ChIP-seq provides only a snapshot of TF activity at the moment of cell lysis and thus may be inefficient at detecting transient or infrequent TF binding events. Finally, while robust for non-cell type-selective, tissue-level analyses, it is often challenging to interpret ChIP-seq data obtained from complex tissues such as the brain, which is comprised of many different interconnected cell types. Because of this limitation, efforts have recently been made to modify ChIP-seq for cell type-specific use, either through physical nuclear sorting^10,13^, or conditional expression and subsequent isolation of tagged nuclei^5,14^ or chromatin-associated enzymes^15^ prior to ChIP. However, these methods thus far have been limited to highly abundant targets, such as histone modifications^5,10^ and transcriptional coactivators^15^, and may be complicated by potential disassociation-related artifacts^16^. Therefore, it is unclear if ChIP-seq is feasible from sorted or isolated nuclei for less abundant TFs.

An alternative approach is to record TF binding events by fusing the TF of interest to DNA-modifying enzymes^17–20^. Prominent among these are two techniques: DNA adenine methylation identification (DamID)^17^, which records TF binding through local adenine methylation by an *E. coli* Dam methylase fused to a TF of interest, and calling cards^18,21^, in which a TF is fused to a transposase enzyme and binding events are recorded through transposon insertion proximal to the TF binding site. Importantly, TF-tagging techniques do not require TF-specific antibodies and have the ability to record and integrate occupancy information across time^22^, while requiring very little starting material^23^. Further, these approaches offer the potential for cell type-specificity through conditional expression of the TF-enzyme fusion protein. In this way, DamID has been successfully implemented for cell type-specific profiling^24^, primarily in *Drosophila*^23^ but also with some studies in cultured mammalian cells^25–27^ and embryos^26^. Meanwhile, calling cards has also been successfully applied to yeast^28^ and mammalian cell^18^ model systems. However, neither of these methodologies has to date been implemented for TF recording in postnatal mammalian model systems, such as mice.

Here we adapt calling cards for *in vivo* use by delivering this system to the mouse brain via adeno-associated virus (AAV). This method, in the mold of traditional calling card technologies^18^, works by first expressing the *hyperPiggyBac* (hypPB) transposase within a cell and providing donor transposons. HypPB inserts donor transposons at TTAA sites throughout the genome, leaving permanent marks, or calling cards, at these loci. These transposons can later be sequenced and mapped to the genome to record the history of hypPB localization across the genome. HypPB-mediated insertions can be used to assess TF binding in two ways: 1) hypPB may be fused to a TF of interest, so that the TF directs the insertion of transposons near its genomic binding sites^18^, or 2) unfused hypPB directly interacts with the bromodomain and extra-terminal domain (BET) protein, BRD4, and directs transposon DNA into BRD4-associated genomic regions^29,30^, most prominently active super enhancers^7^. We establish that calling card systems can be delivered to the mouse brain via AAV and that these components successfully record TF occupancy without the need for a TF-specific antibody. We then create a conditionally-expressed, Cre recombinase-dependent version of AAV-mediated calling cards, termed Flip-Excision, or FLEX, calling cards and demonstrate, as a proof-of-principle, the ability of this system to record cell type-specific TF occupancy profiles in the brain. Lastly, we provide evidence that through continued transposon insertion, FLEX calling cards can record and integrate TF binding events over extended time periods following viral delivery, providing insights into transient TF activity that would be otherwise missed with endpoint measures such as ChIP-seq.

## Results

### Intracranial delivery of calling cards via AAV invokes widespread transposon insertion in the mouse cortex

In order to perform transposon calling cards in mammalian cells, two basic components are required: the hypPB transposase (or a TF-hypPB fusion) and donor transposons^18^. We sought to develop an *in vivo* method to efficiently deliver calling card components throughout the mouse brain to identify TF-associated sites. We first tested AAV as a means for calling card reagent delivery, as viral *piggyBac* delivery methods have been successful in other organ systems previously^31,32^. We packaged hypPB and donor transposons carrying TdTomato reporter genes into separate AAV serotype 9 (AAV9) vectors, which efficiently transduce neuron and astrocyte populations^33^, and intracranially injected these vectors into the cortices of postnatal day 0-1 (P0-1) mice. Animals were sacrificed at P21 for analysis (**Fig 1A**). We analyzed hypPB expression with *in situ* hybridization (**Fig 1SA**) and transposon-derived TdTomato immunofluorescence (**Fig 1B**) across the brain and observed widespread viral transduction in the neocortex, hippocampus, and inner brain structures. As expected with the AAV9 serotype^33^, the vast majority of transduced cell types were neurons and astrocytes (**Fig 1C-D**). These results demonstrate that calling card reagents can be efficiently delivered to the mouse brain by AAV.

**Figure 1.**
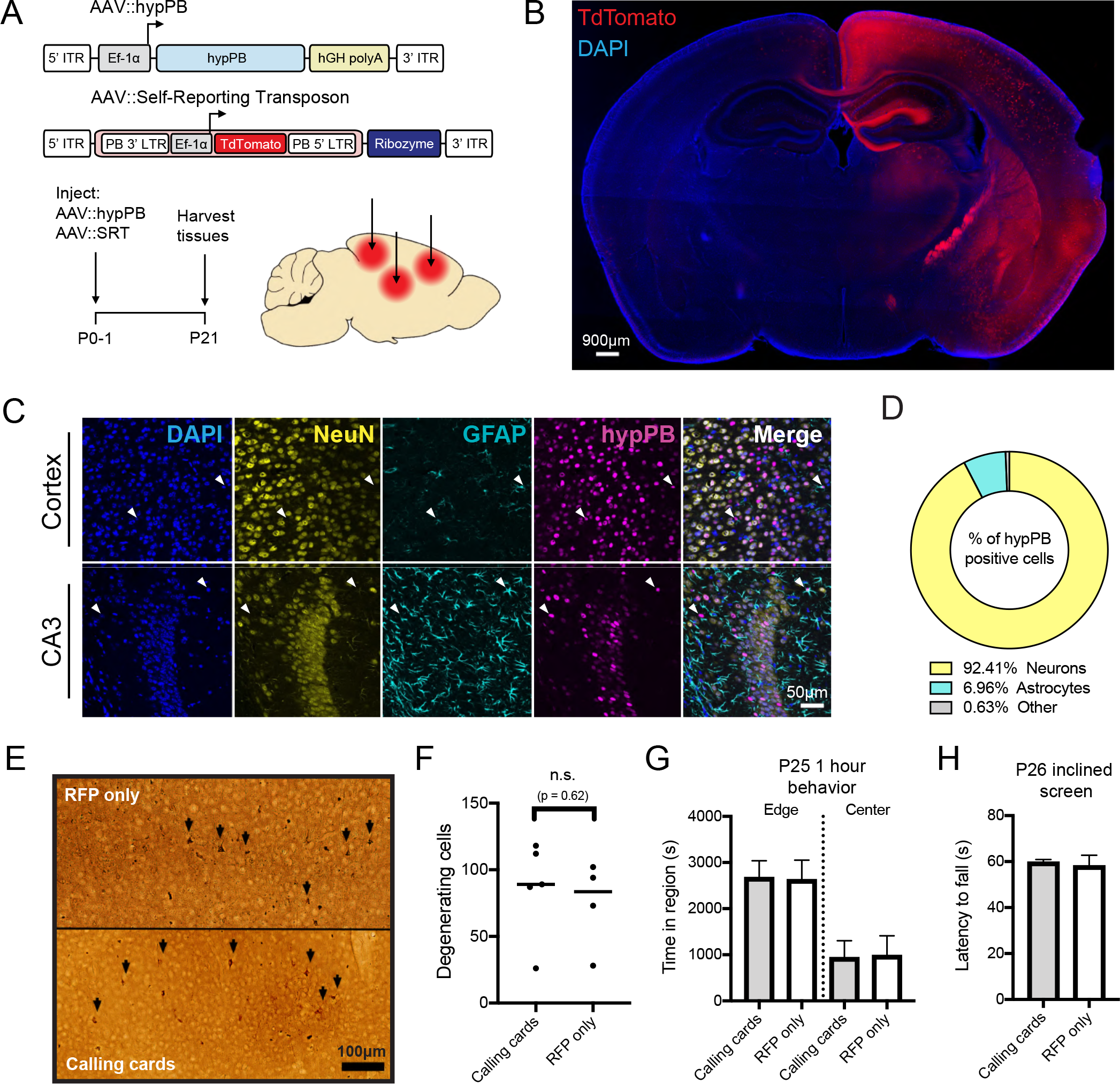
Co-AAV9 intracranial injection efficiently delivers calling cards to the cortex. (A) Experimental paradigm and AAV constructs. Arrows represent approximate AAV injection sites. (B) Coronal section of brain injected unilaterally at P0-1 with AAV∷hypPB and AAV∷SRT, displaying widespread expression of SRT-derived TdTomato fluorescence throughout the brain (C,D) Co-immunofluorescence showing hypPB expression in neurons and astrocytes. (C) Representative images display co-localization of hypPB with neuronal (NeuN) and astrocyte (GFAP) markers in the cortex and hippocampus. Arrowheads show examples of hypPB-positive astrocytes. (D) Majority of hypPB-positive cells transduced with AAV9 are NeuN-positive neurons and GFAP-positive astrocytes (n=1005 myc(+) cells, counted across cortical image fields from 5 mice). (E) Representative images of silver staining in dorsal cortex to screen for degenerating cells (black arrows) in mice intracranially injected at P0-1 with the (top) red fluorescent protein (RFP) or (bottom) calling cards viruses (AAV∷hypPB and AAV∷SRT) and sacrificed at P28. (F) Quantification of silver-positive cells in dorsal cortex revealed injection with either virus produces limited neurotoxicity that did not significantly differ between groups (two-tailed, unpaired Student’s t-test, p > 0.05). (G-H) Mice injected at P0-1 with SRT calling cards (n=21) or control, RFP only (n=24) viruses displayed no significant differences in anxiety related behavior (center/edge dwelling; G) or motor skills (inclined screen test; H) relative to control. See Fig 3S for further behavioral and developmental assessments of these groups. All group comparisons were done with two-tailed, unpaired Student’s t-test, with Bonferroni corrected α=0.05 as a significance threshold (including all tests in Fig 3S).

Earlier implementations of the calling cards method (i.e. BrokenHeart) mapped transposon insertions by directly sequencing genomically-inserted transposon DNA^18,21^ (**Fig 2SA-B**). However, our group recently developed a specialized calling cards donor transposon, termed a “self-reporting transposon” (SRT), which allows for amplification of each insertion via RNA transcription and highly efficient mapping of transposition events through deep sequencing of transposon-derived RNA^34^ (**Fig 2SA,C**). We first sought to directly compare traditional DNA calling cards (BrokenHeart) to RNA calling cards (SRT) in AAV systems. To do this, we intracranially injected P0-1 mice with AAV∷hypPB and either AAV∷BrokenHeart or AAV∷SRT. At P21, we isolated DNA or RNA from cortex samples and generated and sequenced calling cards libraries (**Fig 2SB-C**). We found that SRT reads mapped much more consistently to the mouse genome than BrokenHeart, where the majority of alignment was to the original AAV episomal sequence. We then mapped transposon insertions from these reads. While insertions from the two methods reliably mapped to similar genomic locations (**Fig. 2SF**), we were able recover an order of magnitude greater total number of insertions from animals receiving AAV∷SRT compared to those receiving AAV∷BrokenHeart (**Fig 2SE**). In summary, we found that SRTs provide a much greater sensitivity for recovering insertion events from AAV than traditional DNA methods. Finally, we tested whether SRT calling cards could also be delivered efficiently via adult intraparenchymal stereotactical cortical injection to precise coordinates, as this is more standard for AAV delivery than P0-P1 injection. Indeed, after delivery of AAV∷hypPB and AAV∷SRT to three P107 adult mice (euthanized one month later, at P136), we observed a near-equivalent read mapping rate (**Fig. 2SD**), insertion total (**Fig. 2SE**), and insertion localization (**Fig. 2SF**) as in P0-1 SRT delivery. Thus, AAV calling cards systems are functional *in vivo* and can be delivered to the mouse brain at various timepoints and to targeted locations.

### AAV calling cards delivery to the mouse brain does not induce excess degeneration, weight loss, or behavioral/developmental defects

An important question with all calling card technologies is whether continued transposon insertion is deleterious, particularly after long-term delivery to a living mammalian system such as the mouse brain. To address this question empirically, we intracranially delivered either calling card viruses (AAV∷hypPB/AAV∷SRT) or control (RFP only) viruses to P0-1 mouse pups, and then assessed neuronal degeneration, attainment of developmental milestones, anxiety related behavior, and balance/strength/coordination during the first 4 weeks of life. Four weeks after viral delivery, a small population of degenerating cells were found in the cortex, near the injection site, however there was no significant difference between calling card and RFP-only injected animals, suggesting that the observed degeneration was likely due to needle injury (**Fig 1E-F**). Likewise, no significant differences were observed in weight, righting from back, or edge/center dwelling, indicating that calling cards-injected animals develop normally and display no overt anxiety-like behavior (**Fig 1G** **and Fig 3SA-C**). Finally, in a sensorimotor battery, calling cards-injected animals displayed a largely normal phenotype, with only one test (60° inclined screen climbing) having a significant reduction compared to RFP-only controls. Further, these animals performed normally on the 90° inclined screen and inverted screen tests, which are even more difficult tests of balance and strength than the 60° inclined screen (**Fig 1H** **and Fig 3SD-I**). In summary, AAV SRT calling card reagents did not result in excess degeneration, weight loss, or behavioral/developmental defects, suggesting that genomic transposon insertion does not introduce significant toxicity or deleterious effects to the animal.

### Unfused hypPB delivered with AAV profiles active REs in the brain

BRD4 acts as a “chromatin reader” by binding to acetylated lysine residues on histones^35–37^ and regulating gene transcription^38,39^. Accordingly, BRD4 is strongly associated with active, TF-bound REs, most prominently super enhancers^7,40^. Importantly, BRD4 has a known affinity for the unfused hypPB protein^29^, and consequently unfused hypPB insertions are greatly enriched at known BRD4 binding sites^29^ such as super enhancers^30^. Thus, we aimed to test the hypothesis that unfused hypPB transposon insertion can be used to identify active REs in the brain.

We first analyzed the sensitivity and specificity of unfused hypPB insertions for identification of active super enhancers in a pure, *in vitro* cellular population of neuroblastoma (N2a) cells. To do this, we transfected these cells with the AAV∷hypPB and AAV∷SRT plasmids, from which we collected a total of 806,653 insertions. We then downsampled this library into randomly selected pools of various insertion totals and used peak calling to identify regions of significantly enriched insertion density in each pool, at a range of significance thresholds. Using a previously published N2a H3K27ac ChIP-seq dataset^12^ to independently define active super enhancers in this population, we assayed sensitivity and specificity of calling card insertion peaks for identifying these regions. From this, we observed that calling card peaks are highly specific for active super enhancers across a range of sensitivities, with a high area under the receiver-operator characteristic curve (0.82; **Fig 4SA**). Further, we observed a high sensitivity for super enhancer identification, even at low insertion totals (e.g. sensitivity of up to 0.8 from only 10,000 insertions), that increases steadily with increasing number of insertions (**Fig 4SB**). Thus, unfused hypPB calling card profiles can be used to identify active super enhancers *in vitro*.

We next asked whether AAV∷hypPB could identify active REs, including super enhancers, in the brain. To do this, we combined all 3,732,694 insertions collected from two mice injected with AAV∷hypPB and AAV∷SRT at P0-1 and defined significantly enriched insertion peaks (7,031 peaks; p<10^−30^), which we predict to be BRD4-bound REs. We observed that insertion density at these peaks was highly correlated between the two animal replicates, indicating a high reproducibility of this method (**Fig 2A**). To assess whether insertion peaks represented active REs, we compared our calling card data to ENCODE ChIP-seq datasets^33^ of enhancer-associated histone modifications^4^ in the developing mouse cortex. At the 7,031 significantly enriched insertion peaks, we observed a strong enrichment of active enhancer-associated histone modifications H3K27ac and H3K4me1 and a de-enrichment of the repressive mark H3K27me3 (**Fig 2B-E**). We then used a previously published^41^ P14 H3K27ac ChIP-seq data from mouse cortex to independently define active enhancers and super enhancers and asked whether calling card peaks significantly overlapped these regions. We observed that the majority of insertion peaks intersected with H3K27ac-defined enhancers, significantly higher than when insertion peak coordinates are randomized (**Fig 2F**). Similarly, calling card peak intersection with super enhancers is also significantly higher than chance (**Fig 2G**). As expected for a BRD4-mediated mechanism, unfused hypPB calling card profiles identify only a subset of all enhancers, but do intersect the majority of super enhancers (**Fig 2H-I**). Of note, the reference ChIP-seq dataset used for enhancer and super enhancer identification encompasses REs from all cortical cell types, while calling card peaks are derived only from transduced cells (i.e. neurons and astrocytes), thus likely underrepresenting RE sensitivity. Further, this overlap analysis was performed using our standard, highly rigorous significance threshold for peak calling (p=10^−30^), however we have also performed these analyses at a range of p-value thresholds to confirm the finding is robust to this parameter (**Fig 4SC-D**). Together, these data support that AAV-mediated calling card insertion profiles of unfused hypPB can be used to identify putative REs in the brain.

**Figure 2.**
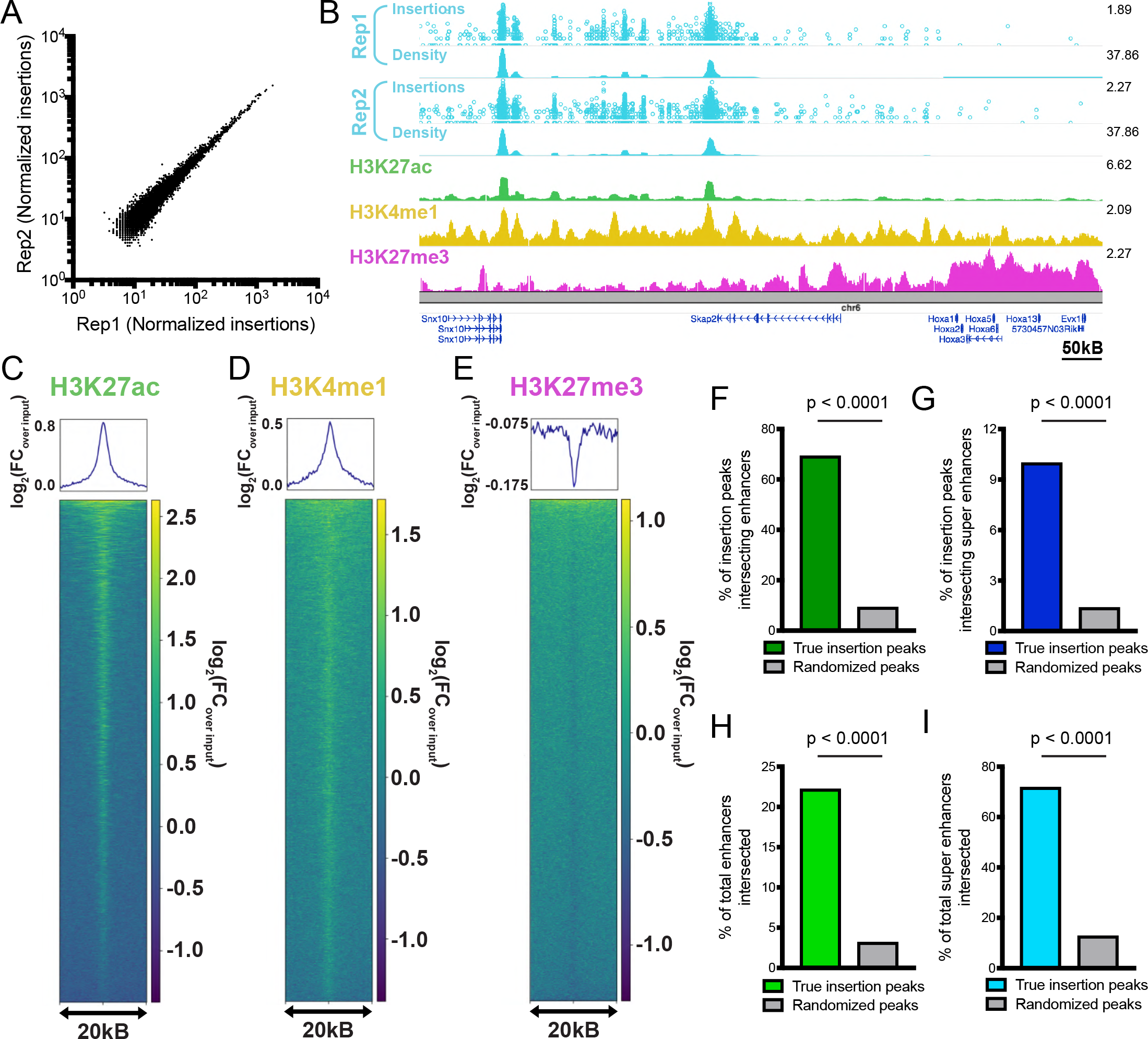
Unfused hypPB-directed calling cards insertions identify active enhancers and super enhancers in the brain. (A) Normalized insertion totals in two littermate C57BL/6J mice (Rep1 and Rep2) at 7031 significantly-enriched insertion peaks (p<10^−30^) displaying high correlation between replicates (R=0.994). (B-E) Unfused hypPB-directed insertions are highly enriched for the active enhancer marks H3K27ac and H3K4me1 and depleted for suppressive mark H3K27me3. Representative image (B), heatmaps, and enrichment plots (C-E) of H3K27ac, H3K4me1, and H3K27me3 density at 7031 significantly-enriched insertion peaks in two littermate mice. In (B), top track of each insertion replicate displays unique insertions, where each circle = 1 unique insertion and y-axis represents number of reads supporting each insertion on a log_10_ scale, and bottom track displays normalized local insertion density across the genome (insertions per million per kB). Y-axis of ChIP-seq data represents read depth with smoothing filter applied. Heatmaps and enrichment plots are centered on insertion peaks and extend 10kB in either direction. Relative enrichment quantifications displayed in log_2_(fold-change over ChIP-seq input). (F,G) Percentage of 7031 significantly-enriched insertion peaks with at least 1 basepair (bp) intersection with a H3K27ac-marked enhancer or super enhancer. Gray bar represents intersections after randomizing genomic coordinates of insertion peaks. χ^2^ test with Yates correction: p<0.0001. (H,I) Percentage of H3K27ac-marked enhancers and super enhancers with at least 1 bp intersection with a significantly-enriched insertion peak. χ^2^ test with Yates correction: p<0.0001.

### FLEX calling cards system allows for cell type-specific profiling of REs in the brain

Gene-based TF-tagging systems such as calling cards offer the potential for cell type-specific profiling through conditional expression of the TF-enzyme fusion. Thus, we generated a Cre-dependent calling cards system, termed FLEX calling cards, and tested the ability of this system to record cell type-specific RE activity or TF binding in complex tissues without isolation of the cell population of interest. In the FLEX system, the reverse complement of the hypPB or TF-hypPB gene is positioned downstream of a strong, ubiquitous promoter and is flanked by two sets of loxP sites. In the presence of Cre, the transposase gene is flipped to the correct orientation and is expressed. To confirm Cre-dependence of the FLEX system, we co-transfected the Cre-dependent hypPB virus, AAV∷hypPB FLEX, into HEK293T cells along with the BrokenHeart reporter plasmid (**Fig 5SA**), which expresses TdTomato only after genomic transposon insertion^42^. We observed BrokenHeart-derived TdTomato fluorescence only in cells that received both the FLEX calling card constructs and a Cre expression plasmid, demonstrating that this system is Cre-dependent (**Fig 5SA**).

As a proof of principle, we focused on two prominent and well-studied Cre-driver mouse lines, Syn1∷Cre and GFAP∷Cre, which direct expression to neurons and astrocytes, respectively. We packaged the AAV∷hypPB FLEX plasmid into the AAV9 vector and intracranially co-injected it along with AAV∷SRT into P0-1 mouse pups of either the Syn1∷Cre or GFAP∷Cre genotype, along with Cre(−) littermates as controls. We euthanized the animals at P28, isolated cortical RNA, and sequenced and mapped insertions across the genome (**Fig 3A**). Immediately apparent upon sacrifice was that brains of Syn1∷Cre positive animals were noticeably more red than their negative littermates, even to the naked eye (**Fig 5SB**), a result of the transposition-enhanced TdTomato reporter expression derived from the AAV∷SRT construct. This change in color was striking for Syn1∷Cre brains, but not as apparent in GFAP∷Cre animals, an observation that is consistent with the relative abundances of transduced neurons and astrocytes (**Fig 1C-D**). In Syn1∷Cre brains, we analyzed TdTomato expression with immunofluorescence and noted a marked increase in neurons of Cre(+) animals but not Cre(−) littermates (**Fig 5SC**). We then sequenced insertions in Cre(+) and Cre(−) littermates from each line and observed a significant increase in insertion events in positive animals as compared to their negative littermates (**Fig 5SD**).

**Figure 3.**
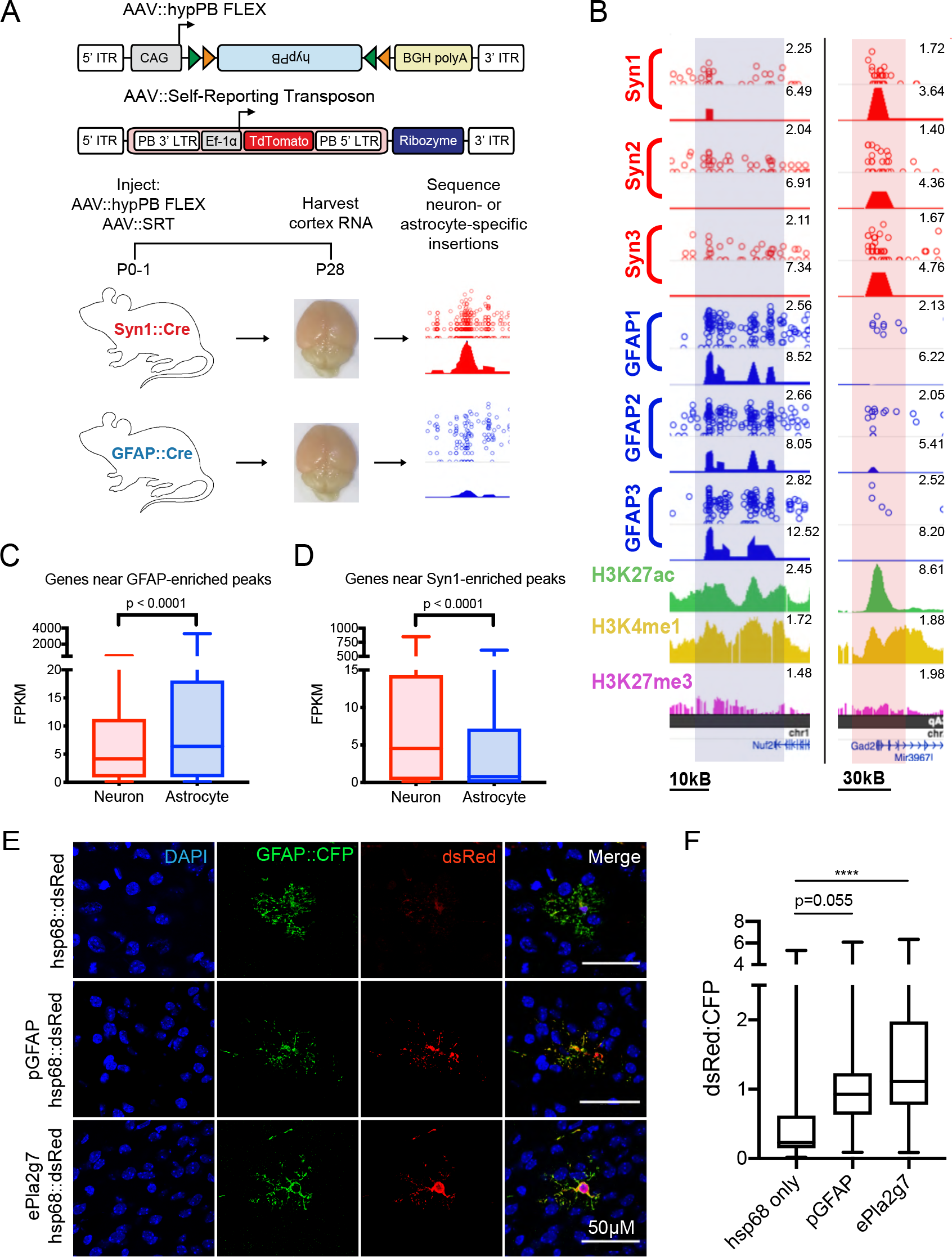
FLEX calling cards system generates cell type-specific RE profiles. (A) AAV constructs and experimental design in Syn1∷Cre and GFAP∷Cre animals. (B) Examples of differentially enriched insertion peaks near genes preferentially expressed in neurons (right) or astrocytes (left). (C,D) Quantifications of neuron and astrocyte specific expression of genes near (C) GFAP∷Cre [Neuron_median_ = 4.16 FPKM, n = 1180 genes; Astrocyte_median_ = 6.38 FPKM, n = 1180 genes] or (D) Syn1∷Cre [Neuron_median_ = 4.55 FPKM, n = 540 genes; Astrocyte_median_ = 0.78 FPKM, n = 540 genes] enriched insertion peaks at a stringent peak-calling significance threshold (p=10^−7^) showing significant preferential expression in the expected cell type (two-tailed Mann-Whitney U test: p<0.0001). (E) A GFAP∷Cre enriched insertion peak proximal to the *Pla2g7* gene (ePla2g7; see Fig 6SE for peak coordinates) was cloned into a plasmid upstream of the hsp68 minimal promoter and a dsRed reporter gene and co-delivered along with a GFAP∷CFP plasmid to ventricle-proximal glia, including astrocytes, with postnatal astrocyte labeling electroporation (PALE)^46^. Expression of dsRed is enhanced by both the canonical GFAP promoter (pGFAP; positive control) and ePla2g7. (F) Quantification of dsRed expression enhancement in CFP(+) astrocytes by pGFAP and ePla2g7. n=34-42 CFP(+) cells from 3 brains per condition (one-way ANOVA with Dunnett’s multiple comparisons test; ****p<0.0001). pGFAP_mean diff._ = 0.66, 95% C.I._diff_ [0.01 : 1.33]. ePla2g7_mean diff._ = 1.22, 95% C.I._diff_ [0.58 : 1.86].

We next sought to test whether FLEX calling cards with unfused hypPB could identify cell type-specific REs. To do this, we identified insertion peaks that were differentially enriched in either Syn1∷Cre over GFAP∷Cre or GFAP∷Cre over Syn1∷Cre by a count-based statistical comparison and asked whether genes near these differentially enriched peaks are more likely to be expressed in neurons or astrocytes, using a previously published and widely used cell type-specific RNA-seq dataset^43^. As predicted, we found that as our significance threshold for defining differentially enriched insertion peaks became more stringent, the RNA expression of nearest gene sets became more cell type-specific (**Fig 5SE-F**). At a stringent significance threshold of p=10^−7^, we compared all nearest genes to Syn1∷Cre or GFAP∷Cre enriched insertion peaks, and found significant differences in astrocyte versus neuron expression in the expected directionalities (**Fig 3C-D** **and Fig 5SG**). Of note, these neuron and astrocyte RNA enrichments were observed despite using proximity as a means for enhancer-gene pairing, which while widely used for analyses such as these^3^, is likely only identifying the correct gene of interest in a subset of pairs^6^. Lastly, we inputted these gene sets into an unbiased cell type-identification tool (CSEA^44^) and successfully identified cortical astrocyte and neuron populations for genes near GFAP∷Cre and Syn1∷Cre enriched insertion peaks, respectively (**Fig 5SH-I**). Together, these data indicate that peaks derived from FLEX calling cards insertion profiles recorded by unfused hypPB represent cell type-specific REs responsible for driving expression of cell type-enriched genes.

Lastly, we sought to functionally validate the enhancer activity of a subset of the novel astrocyte-enriched REs by testing whether these regions could enhance the expression of a dsRed reporter gene in astrocytes *in vivo*. We chose 4 candidate astrocyte-enriched REs, based on their size, cell type-specific activity, and astrocyte/neuron RNA expression of their nearest genes (**Fig 6SA-E**). We then cloned these candidate REs upstream of the hsp68 minimal promoter driving a dsRed reporter gene. As a positive control, we also cloned the canonical GFAP promoter (pGFAP)^45^ into the same location upstream of hsp68∷dsRed. To test the functional enhancer activity of these REs *in vivo*, we delivered these plasmids, along with a separate plasmid carrying a CFP reporter under the GFAP promoter for astrocyte identification, via postnatal astrocyte labeling electroporation (PALE)^46^. At P7, mice were euthanized and brains were collected for immunohistochemistry. This method specifically targeted astrocytes in the cerebral cortex, noting that >96% of dsRed(+) cells in this region were also GFAP∷CFP(+) (**Fig 6SG**). As expected, the positive control (pGFAP hsp68∷dsRed) plasmid exhibited enhanced dsRed fluorescence in astrocytes, relative to a negative control plasmid carrying only hsp68∷dsRed, that approached statistical significance (p=0.055; **Fig 3E-F** **and Fig 6SF,H**). We then quantified enhancer activity of our candidate REs and observed a significant enhancement of dsRed fluorescence for three of the four candidates (**Fig 3E-F** **and Fig 6SF,H**). Thus, these astrocyte-enriched REs display functional enhancer activity in astrocytes in the mouse brain at P7. Next, we repeated this experiment allowing mice to age to P21 to allow further astrocyte maturation prior to euthanasia. To our surprise, at this timepoint we observed a change in localization of dsRed expression in brains receiving the minimal promoter hsp68∷dsRed construct, with fewer GFAP∷CFP(+) astrocytes and a new population of NeuN(+) neurons labeled with dsRed in the cortex (**Fig 7SA-B**). This suggests that the PALE method does deliver plasmids to neurons or neural progenitors in addition to astrocyte progenitors, but that expression via the hsp68 promoter in neurons does not arise until later in postnatal development. Strikingly however, in animals that received pGFAP hsp68∷dsRed or any of the RE candidate plasmids, dsRed expression was contained to GFAP∷CFP(+) astrocytes, suggesting that in addition to enhancing expression in astrocytes, these RE sequences also repress activity of hsp68 in neurons (**Fig 7SA-B**). Indeed, even eMms22l, which was not yet active at P7, displays this functional enhancer phenotype at P21. Overall, these data demonstrate that cell type-specific REs derived from FLEX calling cards functionally regulate cell type-specific gene expression in the brain.

### Fusion of hypPB to the promoter-binding transcription factor SP1 records SP1 occupancy

A key feature of calling cards is the ability to record binding of a TF of interest using TF-hypPB fusions. To demonstrate calling card TF recording *in vivo*, we fused hypPB to a sequence-specific DNA binding general TF, SP1, which binds to gene promoters and is involved in transcription^47,48^, and cloned this fusion gene into the FLEX calling cards system for cell type-specific use (**Fig 4A**). As full length SP1 is too large to be efficiently packaged into AAV, we instead used a truncated version of SP1 containing the C-terminal 621 amino acids, which includes the DNA-binding domain and has been shown to be sufficient to replicate sequence-specific binding of full length SP1^49^. To test this system, we intracranially co-injected AAV∷SP1(621C)-hypPB FLEX along with AAV∷SRT into P0-1 mice of the Syn1∷Cre line, sacrificed animals at P28, generated and sequenced SRT libraries from cortical RNA samples, and compared insertion profiles to that of unfused hypPB. Consistent with the affinity of SP1 for proximal promoters, we found that insertions were significantly enriched upstream of TSS, as compared to unfused hypPB insertion profiles (**Fig 4C**). Next, we defined differentially enriched insertion peaks in SP1(621C)-hypPB profiles over unfused hypPB (p<10^−15^) and found that the majority of significant enrichments occur in gene promoters (**Fig 4D-E**). Finally, at these SP1 peaks, we performed motif discovery and were able to identify enrichment of the canonical SP1 binding motif, GGGCGGGG^18^ (**Fig 4F**). Thus, fusion of SP1 to hypPB and delivery via AAV identifies SP1 binding sites in the mouse brain.

**Figure 4.**
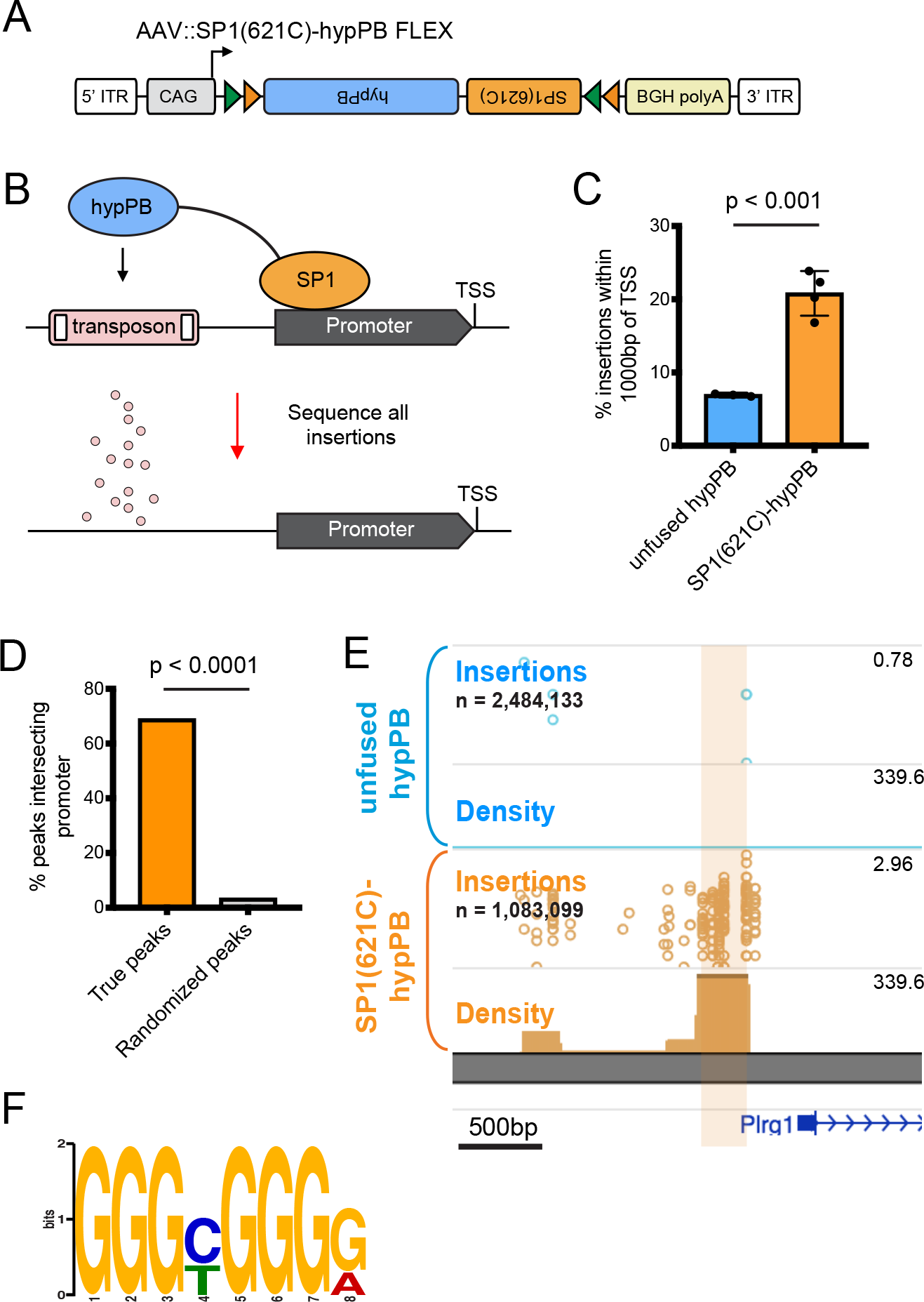
Fusion of SP1 to hypPB in FLEX calling cards system records SP1 occupancy. (A) Schematic of AAV∷SP1(621C)-hypPB FLEX construct. (B) Fusion of the promoter-binding TF SP1 to hypPB directs insertions to promoter-proximal TTAA sites. (C) Percentage of total insertions within 1000 basepairs (bp) of a transcription start site (TSS), displaying increased promoter-directed insertions upon SP1 fusion as compared to unfused hypPB (n = 3-4 mice per group; two-tailed, unpaired Student’s t-test: p<0.001, t=7.66, df=5, 95% C.I._diff_ [9.22 : 18.54]; unfused hypPB_mean_ = 6.9%, SP1(621C)-hypPB_mean_ = 20.8%). Error bars represent SD. Total SP1(621C)-hypPB insertions: 1,083,099. Total unfused hypPB insertions: 2,484,133. (D) Percentage of significant SP1 insertion peaks differentially enriched over unfused hypPB (p<10^−15^; 1596 intersecting out of 2316 total peaks) intersecting promoter-proximal regions (1000bp on either side of TSS) compared to randomized peak coordinates (78/2316). χ^2^ test with Yates correction: p<0.0001. (E) Representative insertion peak displaying significantly increased insertion density near the TSS of the *Plrg1* gene. (F) Highest information content motif in the sequences flanking the center of significantly enriched SP1(621C)-hypPB insertion peaks (p<10^−15^) is the canonical SP1 binding motif (GGGCGGGG; p<10^−42^).

### FLEX calling cards provides historical TF binding information through longitudinal TF recording

An intriguing potential use of calling card technologies is in the recording of TF binding over an integrated period of time. Such a method, which is not possible with endpoint TF profiling methods such as ChIP-seq or CUT&Tag, could empower novel studies in which historical TF binding information would be useful, such as during cellular development or differentiation. Further, by integrating signal over time, longitudinal calling cards may report transient binding events which would be otherwise missed with endpoint-only measures.

To test whether FLEX calling cards could report integrated, historical TF occupancy, we asked whether we could recover transient SP1 promoter binding events and successfully read them out at a later date. Importantly, consistent with the known role of SP1 in regulating gene expression^18,47,48^, we observed that expression of genes genome-wide was on average correlated with the number of SP1-directed promoter insertions (**Fig 5A-B**). Thus, we predicted that should a gene be expressed early, but not late, in the lifetime of the animal, this transient event could be marked by SP1 binding and be recoverable via SP1 calling cards at a later timepoint.

**Figure 5.**
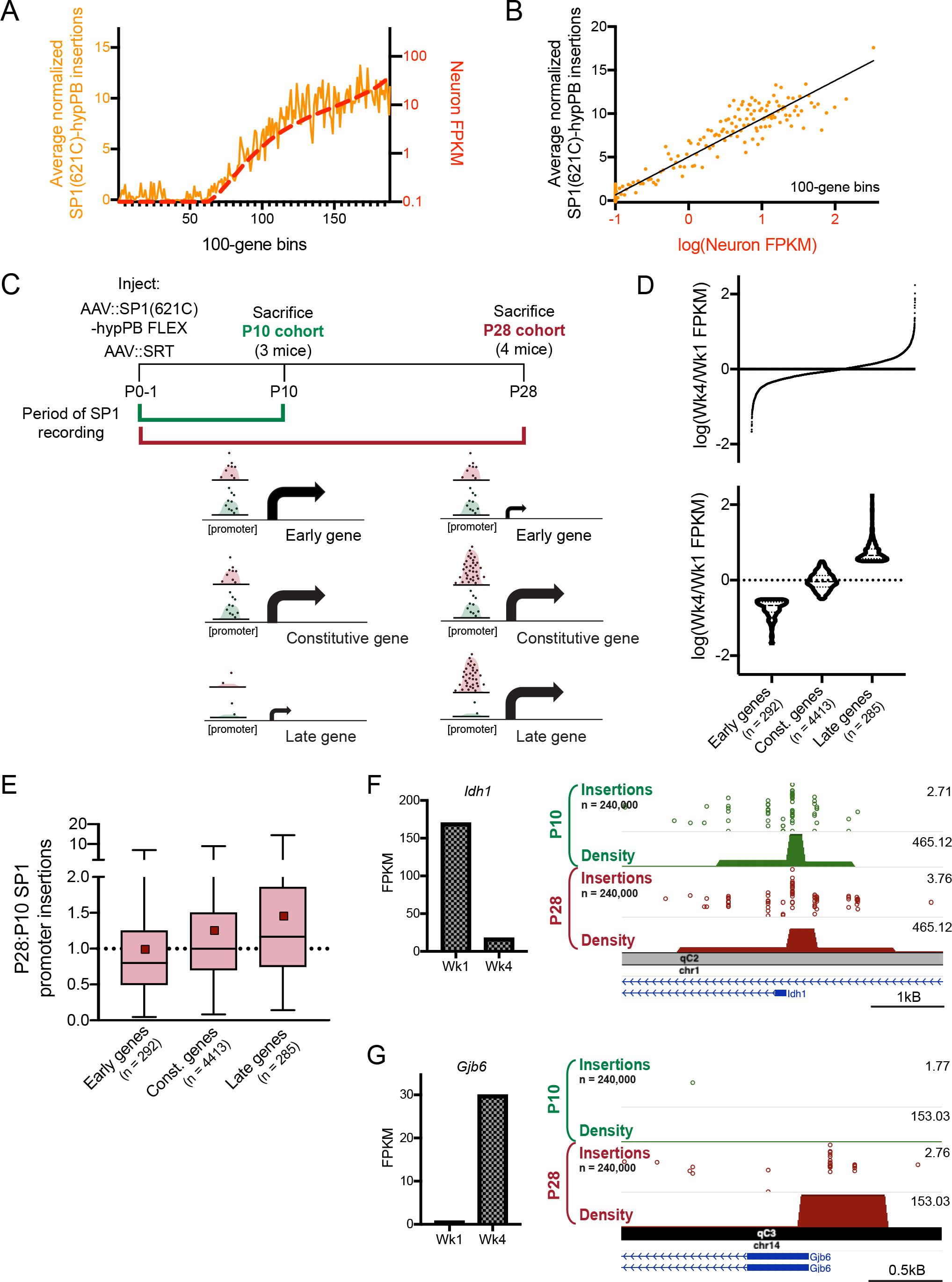
Longitudinal SP1 profiling reports integrated record of SP1 binding. (A-B) Normalized number of SP1(621C)-hypPB directed insertions at promoter proximal regions after subtraction of unfused hypPB insertions, versus neuron-specific gene expression for all genes, binned and averaged into 100-gene bins. In (A), left y-axis represents number of promoter insertions normalized to 10^6^ total insertions in the sample and right y-axis displays neuron-specific RNA expression from Zhang et al., *Journal of Neuroscience*, 2014. (B) displays strong correlation of SP1(621C)-hypPB promoter insertions with gene expression after subtraction of unfused hypPB insertions (R=0.96, p<0.0001). (C) Experimental paradigm and predicted temporal SP1 occupancy for early, constitutive, and late expressing genes. (D) (top) Distribution of Wk4/Wk1 expression ratios for all expressed and SP1-bound genes and (bottom) categorization into “early” (log(Wk4/Wk1 FPKM) < −0.5), “constitutive” (−0.5 < log(Wk4/Wk1 FPKM) < 0.5), and “late” (log(Wk4/Wk1 FPKM) > 0.5) gene sets. RNA-seq from Lister et al., *Science*, 2013. (E) SP1-derived promoter insertions for early, constitutive, and late gene sets, demonstrating efficient capture of transient SP1 binding events at early gene promoters and continued integration of constitutive and late gene promoters in the P28 cohort relative to the P10 cohort (One-way ANOVA [F(2, 4987) = 16.92], p<0.0001). Early genes_mean_ = 0.98, const. genes_mean_ = 1.25, late genes_mean_ = 1.45. Red square = mean, line = median. (F) Example of early expressed gene (*Idh1*) displaying equivalent SP1 binding in both cohorts. (G) Example of late expressed gene (*Gjb6*) displaying SP1 binding only in the P28 cohort.

To test this hypothesis, we intracranially co-injected AAV∷SP1(621C)-hypPB and AAV∷SRT into two separate cohorts of P0-1 mice. The first cohort was euthanized at P10, while the second cohort was allowed to continue to record SP1 occupancy until P28 (**Fig 5C**). This time period of postnatal brain development involves several key neurodevelopmental processes^50^, including substantial hippocampal neurogenesis^51^ as well as glial and synaptic maturation^50^, development of the extracellular matrix^50^, and closing of critical periods^50^, which are accompanied by numerous changes in gene and protein expression^52^. For these analyses, we utilized a previously-published cortical RNA-seq dataset^53^ with postnatal timepoints of 1 week (Wk1; ~P7) and 4 weeks (Wk4; ~P28). From these expression data, we then derived and tested three separate predictions (**Fig 5C**). First, for genes expressing at Wk1 but not Wk4, (i.e. “early genes”) we would observe near-equivalent SP1 binding at promoters in both cohorts. Second, for constitutive genes that express equally at Wk1 and Wk4, we would observe continued integration of SP1 binding in the P28 cohort, resulting in increased SP1 insertion density at promoters. And third, for genes expressing only at Wk4 (i.e. “late genes”), we would observe SP1 promoter binding only in the P28 cohort. We defined early genes as having a log(Wk4/Wk1 FPKM) of less than −0.5 (n = 292), late genes as greater than 0.5 (n = 285), and all genes in the middle as constitutive (n = 4413; **Fig 5D**). Indeed, at the promoters of these gene sets, we observed SP1 promoter occupancy to be consistent with our three predictions (**Fig 5E-G**). Importantly, at the promoters of early genes, we observed near-equivalent binding in the P10 and P28 cohorts (mean P28:P10 SP1 promoter insertion ratio = 0.98), despite these genes only being transiently expressed; thus, this system is capable of permanently recording transient TF binding events for retrospective read out at a later date. Together, these data support that TF-hypPB fusions integrate signal over time and provide a historical, integrated picture of TF occupancy.

## Discussion

In this report, we have successfully developed an *in vivo*, virally mediated calling cards approach, for which we have demonstrated utility for TF and RE profiling in the live mouse brain. This technology builds on previously developed *in vitro* calling card methodologies, adapting this method for investigation of epigenetic regulation in the mammalian brain. Further, in proof-of-principle experiments, we demonstrated effectiveness of this protocol for 1) profiling TFs without an antibody, 2) cell type-specific RE profiling without cellular population purification, and 3) integrative recording of TF binding events over time. Our use of SRTs in this paradigm now also enable calling cards for single cell analyses^34^, which expands and highlights the versatility of this toolkit in future studies.

Calling cards technologies, as gene-based systems, have several unique features that provide advantages over biochemical TF profiling methods such as ChIP-seq for certain applications^18,21,54^. One such property of the FLEX version of the AAV calling card methodology is that there is no requirement for physical isolation of cell types or nuclei for cell type-specific analyses. This allows for the same protocol to be used for any cell type of interest, the identify of which is determined by the Cre-driver mouse line used, and avoids potential disassociation-related artifacts^16^. Secondly, while not explored here, one could envision simple manipulations of the AAV calling cards system to allow for temporal control of the system^43^. Such adaptations could allow for innovative studies in which TF binding is recorded only during defined windows of time^22^. In a similar vein, we have demonstrated here the ability of AAV calling cards to integrate TF binding information over time, which will allow for retrospective analysis of historical TF activity in cells. By applying this unique utility to SP1, we identified promoter regions of genes with accumulating SP1 binding across postnatal development and captured transient SP1 binding events at early-expressed genes. Finally, AAV calling cards does not require a TF-specific antibody, allowing for TF profiling for, in theory, any packagable TF, simply by fusing it to hypPB. This being a virally-mediated system allows for simple and rapid application to animal models without the need for expensive and time-consuming breeding. Intracranial injection for a standard size litter of mice can be completed in under an hour. Finally, the non-Cre dependent versions of the system should be equally applicable in other species of interest such as rats and primates. Reagents, cloning strategies, and user-friendly analysis pipelines are available upon request, making AAV calling cards readily available for neuroscience research.

Of course, there are caveats to be considered as well. Most notably, there is potential for induced mutation, given the tendency for transposons to insert into or near critical gene regulatory regions. Indeed, transposon technologies are often used in mutagenesis screens in which transposon-mediated gene disruption can be deleterious^55^. However, in such studies, the transposons are specifically engineered with splice-site gene or enhancer traps, while the SRT used in AAV calling cards only drives expression of a reporter gene and the genomic sequence immediately downstream of its insertion site. Consistent with this, we observed no excess degeneration or behavioral/developmental deficits in AAV calling card-injected animals beyond that induced by needle injury. Further, the transposition rate of the *piggyBac* transposase is inherently low (<20 per cell^56^), suggesting that it is highly unlikely for insertions to disrupt regulatory regions on both alleles in the same cell, and in general, calling card technologies have not exhibited marked deleterious effects in previous reports^18,21^. Nevertheless, it remains possible that a subset of calling cards transposition events could perturb nearby gene expression in a subset of cases. Second, while useful for profiling enhancers and super enhancers more broadly, the natural affinity of hypPB for BRD4 does necessitate that any experiment using a TF-hypPB be accompanied by a control with unfused hypPB only, such that TF binding peaks can be identified with differential peak calling, as was done for SP1 in this report. Future versions of AAV calling cards systems may be improved by fusion of TFs to other, non-BRD4-biased transposases^30^, though the efficiency of transposon re-direction would need to be tested empirically in each case. Finally, it is important to recognize that while we do see a clear Cre-induction of the FLEX calling card system, providing a proof of principle for its use in cell type-specific profiling, we also observed some background insertion events in the absence of Cre, which could be limiting for profiling of rare cell types in which signal is likely to be reduced. We expect future applications of the viral FLEX calling card method to improve as conditional AAV expression systems with tighter control of gene expression become available. Alternatively, simply changing the viral serotype^33^ or promoter^57,58^ could allow for similar analyses in cell types not explored here without the need for Cre-dependent conditional expression.

In summary, we have introduced AAV-mediated FLEX calling cards as a viable method for recording TF binding and active REs *in vivo* and demonstrated its effectiveness in profiling cell type-specific and historical TF and RE activity in the brain. Future applications of this technology to animal models of development and disease could unlock important insights into epigenetic gene regulation in a variety of neuroscience disciplines.

## Acknowledgments

We thank the Genome Technology Access Center in the Department of Genetics at Washington University School of Medicine for genomic analysis. The Center is partially supported by NCI Cancer Center Support Grant #P30 CA91842 to the Siteman Cancer Center and by ICTS/CTSA Grant# UL1 TR000448 from the National Center for Research Resources (NCRR), a component of the National Institutes of Health (NIH), and NIH Roadmap for Medical Research. This publication is solely the responsibility of the authors and does not necessarily represent the official view of NCRR or NIH. We thank the Edison Family Center for Genome Sciences & Systems Biology, specifically Jessica Hoisington-Lopez and MariaLynn Crosby, for assistance with genomic analysis. We thank Bernard Mulvey for providing hsp68∷dsRed plasmids. Finally, we thank the Kevin Noguchi, Brant Swiney, and the Intellectual and Developmental Disabilities Research Center (IDDRC) at Washington University (NICHD U54-HD087011) for neuropathological assessments.

## Author Contributions

Study designed by AJC, AM, TL, JC, MJV, RDM, and JDD. AJC, AM, JC, JH, XC, MNW, and MH designed and generated DNA constructs. AM developed SRT analysis pipelines. AJC, TL, JC, MJV, MS, KM, XC, and RDM generated and analyzed data. Manuscript written by AJC and edited by AM, TL, JC, MJV, TMM, RDM, and JDD.

## Methods

### Animals

All animal practices and procedures were approved by the Washington University in St. Louis Institutional Animal Care and Use Committee (IACUC) in accordance with National Institutes of Health (NIH) guidelines. Transgenic mouse strains used in this study include Synapsin 1 (Syn1)∷Cre (RRID:IMSR_JAX:003966) and glial fibrillary acidic protein (GFAP)∷Cre (RRID:IMSR_JAX:024098). All mice were bred to the C57BL/6J background, with the exception of animals used for adult stereotactic injections, which were wild-type animals of the FVB/N6 background (RRID:IMSR_JAX:001800). At indicated endpoints, mice were anesthetized with isoflurane and perfused with 15ml of cold saline (PBS) prior to tissue collection. Animals in the BRD4 Syn1∷Cre, BRD4 GFAP∷Cre, and P28 SP1 Syn1∷Cre cohorts received pentylenetrazole-induced seizures immediately prior to sacrifice. Unless otherwise noted, brains were either dissected and flash frozen in liquid nitrogen (for molecular analyses) or fixed in 4% paraformaldehyde 24-48 hours, exchanged into 30% sucrose, and either directly frozen at −80°C or cryoprotected in O.C.T. (for immunofluorescence).

To collect newborn litters, pregnant females were monitored daily until giving birth. Newborn pups were injected within 24 hours of being found in the cage, i.e. 0-48 hours after birth; thus, these injections are labeled “P0-1.”

### Cell culture and transfections

HEK293T and N2a cells used in this study were cultured in 1X DMEM with 10% fetal bovine serum (FBS) and grown under standard conditions (37°C; 5% CO_2_). Plasmid transfections in HEK293T cells were carried out with Fugene® 6 Transfection Reagent (Promega, Madison, WI, USA) with the manufacturer’s protocol. Calling cards constructs were delivered to N2a cells via either Fugene 6 or Neon Electroporation (ThermoFisher #MPK10025) with the following settings: 1050V, 30ms, 2 pulses.

### Immunofluorescence and imaging

10μm-thick (for co-localization studies) or 40μm-thick (for imaging of AAV∷BrokenHeart) fixed-frozen sagittal or coronal brain sections were washed with PBS and permeabilized with 0.1% Triton X-100 (Sigma-Aldrich, St. Louis, MO, USA). Non-specific binding was blocked with 5% normal donkey (Jackson ImmunoResearch, West Grove, PA, USA) or goat (Vector Laboratories, Burlingame, CA, USA) serum for 30-60 minutes at room temperature. After blocking, slides were exposed to primary antibody overnight at 4°C, washed three times with PBS, and then incubated with secondary antibodies for 1 hour at room temperature. Nuclei were counterstained with DAPI (Sigma-Aldrich, St. Louis, MO, USA) and coverslips were applied with ProLong Gold Antifade (ThermoFisher, Waltham, MA, USA) or Fluoromount-G (SouthernBiotech, Birmingham, AL, USA) mounting media. Immunofluorescent images of brain sections were acquired with a Nikon A1Rsi or Zeiss LSM 700 confocal microscope and imported into ImageJ (v. 1.51s) for manual cell counts and quantification. For analyses of hypPB expression in various cell types, 5 mice were used, and co-localization was quantified in 2 cortical images from a single section per animal. Antibodies used for immunostaining included chicken anti-GFP (Aves Labs GFP-1020) at 1:1000 dilution, mouse rabbit anti-RFP at 1:400 or 1:500 dilution (Rockland 600-401-379), anti-NeuN at 1:100 dilution (Millipore-Sigma MAB377), rabbit anti-cMyc at 1:250 dilution (Sigma C3956), and goat anti-GFAP at 1:500 dilution (Abcam ab53554).

Cells transfected with AAV∷hypPB FLEX for testing Cre-dependence were live imaged for TdTomato on a Leica DMI 3000B tissue culture microscope. All images were acquired with equal conditions and exposure times for direct comparison.

### In situ hybridization and imaging

10μM-thick, 4% paraformaldehyde fixed-frozen sections were cut and slide-mounted. mRNA encoding for *hyperPiggyBac* (VF1-20268-01) was detected using a custom probe-set designed by Affymetrix (now ThermoFisher) using the Affymetrix ViewRNA ISH Tissue 1-Plex kit (ThermoFisher, QVT0050) and chromogenic signal amplification kit (ThermoFisher VT0200) with the following modifications: Slides were immersed in 4% paraformaldehyde overnight at 4°C prior to *in situ* hybridization, then the baking, deparaffinization, and heat pretreatment steps were omitted (steps 1-3, 5) because sections were not embedded in paraffin. Slides were hybridized either with anti-hypPB or no-probe controls. Following the *in situ* labeling protocol, sections were labeled for 5 minutes with DAPI (1:20,000, Sigma D9542), washed with PBS, and a drop of prolong gold (ThermoFisher P36934) was added while applying the coverslip. Slides were then imaged at 20× magnification on a Zeiss LSM 700 confocal microscope using a Cy3 filterset to detect FastRed fluorescence.

### Analysis of brain degeneration following viral injection

Mice were exposed to either the calling cards viruses (AAV∷hypPB + AAV∷SRT) or RFP-only virus (as a control) via P0-1 injection and sacrificed 28 days later for silver staining (a marker for cells irreversibly committed to cell death). Briefly, animals were heavily sedated and perfused with TRIS fixative in 4% paraformaldehyde followed by vibratome sectioning of brains at 75μM in the coronal plane. Every eighth section across the rostrocaudal extent of the brain was then silver stained as described previously^59^. The number of degenerating neurons was quantified for each animal by a rater blind to treatment who counted the total number of silver positive neurons in the dorsal cortex of each section.

### Mouse behavior, developmental milestones, and sensorimotor battery

Mouse behavior and development was monitored and compared between animals injected calling cards AAV reagents (AAV∷hypPB and AAV∷SRT) or AAV∷RFP only. In addition to weight, which was measured at P8, P14, P25, tests were administered to assess attainment of developmental milestones (P14 righting from back), anxiety related behavior (P25 1 hour behavior, recording time spent in the edge or center of cage), and balance/strength/coordination (P25/26 sensorimotor battery). Procedures were done as previously described^60,61^, with two trials per animal. A break was allowed after the completion of the first set of test trials to avoid exhaustion effects, and the test order was reversed for the second trial for all animals. Walking initiation, ledge performance, platform performance, and pole performance were all administered at P25, while 60° inclined screen test, 90° inclined screen test, and inverted screen test were administered on P26. Prior to testing, mice were given a routine health check, and from this, two animals were excluded; one for runtiness and one for severe hydrocephaly likely derived from needle stick. Further, one litter of RFP-only animals was not able to complete all timepoints (cage flooding) and was thus also excluded. In total, 21 animals were included in the calling card group (11M/10F) and 24 in the RFP-only group (10M/14F), all of which were used for downstream analyses. Test administrators were blinded from treatment group identity during all testing.

### Virus generation and injections

Transposase and donor transposon constructs were cloned into Cre-dependent (FLEX) or Cre-independent AAV transfer vectors and used for *in vitro* transfection or viral packaging. Plasmids were packaged into AAV by the Hope Center Viral Vectors Core at Washington University School of Medicine. For *in vivo* experiments involving P0-1 delivery, transposase and donor transposon viruses (mixed equally by volume) or undiluted RFP-only virus were intracranially injected into the cortex of postnatal day 0-1 (P0-1; 3 sites per hemisphere, 1μl viral mix per site). For adult injections, viruses were delivered to P107 animals intraparenchymally with stereotactic surgery, as previously described^62^. Two sites were chosen for direct, unilateral cortical injection with coordinates relative to bregma of 1.25mm rostral; 1.5mm lateral; 0.55mm depth; and 1.06mm caudal; 1.5mm lateral; 0.55mm depth. 2μl of viral mix was delivered at a rate of 0.2μl/minute.

Viral titers (viral genomes per milliliter) were as follows:

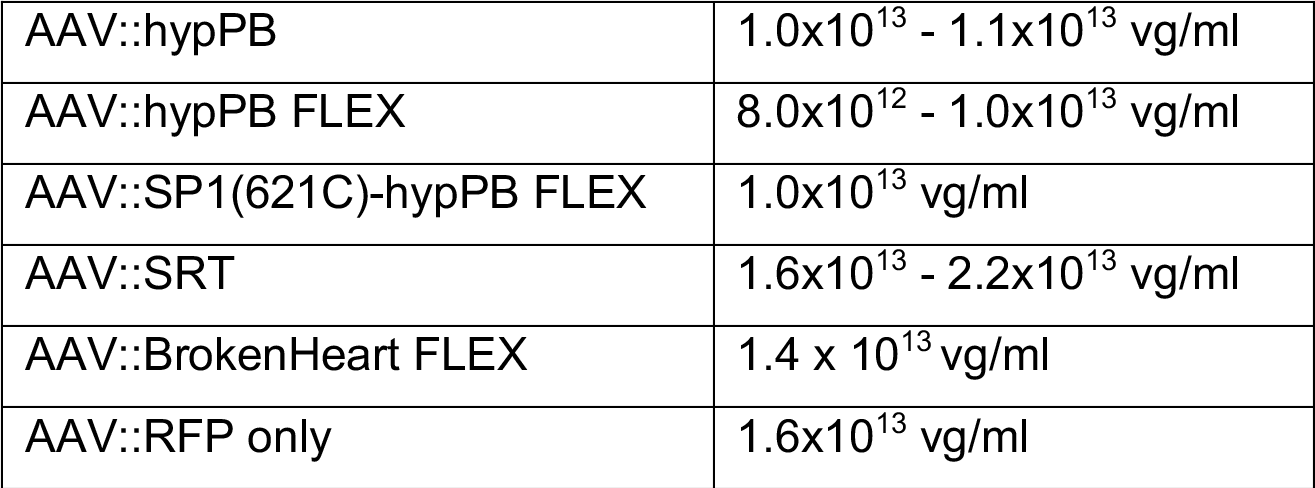

|  |  |
| --- | --- |
| AAV::hypPB | 1.0x10<sup>13</sup> - 1.1x10<sup>13</sup> vg/ml |
| AAV::hypPB FLEX | 8.0x10<sup>12</sup> - 1.0x10<sup>13</sup> vg/ml |
| AAV::SP1(621C)-hypPB FLEX | 1.0x10<sup>13</sup> vg/ml |
| AAV::SRT | 1.6x10<sup>13</sup> - 2.2x10<sup>13</sup> vg/ml |
| AAV::BrokenHeart FLEX | 1.4 x 10<sup>13</sup> vg/ml |
| AAV::RFP only | 1.6x10<sup>13</sup> vg/ml |

### SRT and BrokenHeart library preparation, sequencing, and mapping

SRT libraries were prepared from cortex RNA samples. Prior to library preparation, cortex samples were dissected into 10 separate pieces, from which RNA was independently isolated with the manufacturer’s protocol (Qiagen RNeasy kit, Germantown, MD, USA). This allows for identification of up to 10 independent insertion events into any TTAA site, given that these insertions occur in spatially separate samples. From these RNA samples, transposon sequencing libraries were generated with our bulk SRT protocol^34^. In brief, RNA samples were first reverse transcribed, from which self-reporting transcripts, including flanking genomic sequences, were amplified via PCR. These amplicons were then tagged with universal Illumina sequencing overhangs, with separate indexes for libraries from each dissected piece, allowing for 10 ‘barcodes’ per sample and sequenced on Illumina HiSeq 2500, NextSeq 500, or MiniSeq platforms (Illumina, San Diego, CA, USA).

BrokenHeart libraries were prepared from cortical DNA samples, as previously described^18^. A set of 20 individually barcoded BrokenHeart transposons were pooled and packaged into AAV. Thus, independent insertions into the same TTAA can be uniquely identified via barcode, removing the need to dissect and process tissue samples in separate pools as with SRT. Extracted DNA was self-ligated, amplified with inverse PCR, and sequenced with the Illumina NextSeq 500 platform (Illumina, San Diego, CA, USA).

Sequencing reads obtained from SRT and BrokenHeart libraries were stringently filtered for features of true insertion events (presence of *piggyBac* terminal repeat sequence; intact sequencing adapters, barcodes, and indexes; and a TTAA site) and mapped to build version mm10 of the mouse genome with Novoalign 3 (Novocraft Technologies) (for SRT) or Bowtie2^63^ (for BrokenHeart). Reads aligning to the same TTAA with separate barcodes were considered unique insertions, and all analyses in this report considered all unique insertions equally, independent of read depth. We used BEDtools intersect to count the number of insertions directed to introns, exons, 3’- and 5’-UTRs, and intergenic regions, using annotations from the HOMER package^64^ (**Fig 2SE**).

### Significant insertion peak calling and motif discovery

Significantly enriched insertion peaks were identified via a count-based statistical comparison as previously described^34^. In brief, this pipeline first segments the genome into blocks of constant insertion density. For each block, it calculates the p-value of insertion enrichment relative to a background model assuming uniformly distributed insertions. A user-defined significance threshold is defined, and all blocks surpassing this threshold are considered “significantly enriched insertion peaks”. This “background-free” method for unbiased identification of all significantly enriched genomic regions in a single experimental sample, used here in **Fig 2**, is expected to identify all BRD4-bound regions within the parameters of the calling cards system.

Alternatively, we can define differentially-bound regions between two experimental samples, as was done in **Fig 3, Fig5S, and Fig 6S** for astrocyte- or neuron-specific BRD4 peaks (using the combined insertion pools from three GFAP∷Cre [n = 3,072,163 insertions] and three Syn1∷Cre animals [n = 2,484,133 insertions]) and **Fig 4** for SP1 peaks over unfused hypPB. In this analysis, the pipeline again segments the genome into blocks, but then assigns a p-value to each block based on the differential enrichment between the two samples. As with the background-free pipeline, a user defined p-value threshold is chosen, below which all blocks are considered significantly enriched. Because of the natural BRD4-bias of hypPB, using this differential peak caller allows for identification of binding sites of specific TFs by selecting genomic locations with re-directed insertion density in a TF-hypPB profile compared to an unfused hypPB background.

TF motifs were identified with MEME-ChIP v4.11.2 motif discovery software^65,66^ with -zoops -meme-minw 6 -ccut 250.

### Defining enhancers and super enhancers

Since H3K27ac is a known marker of active enhancers^4,67^ and super enhancers^7,40^, we utilized published P14 mouse cortex^41^ and N2a^12^ H3K27ac ChIP-Seq datasets to define cortical and N2a enhancers super enhancers, respectively. As previously described^7,40^, we used the rank ordering of super enhancers (ROSE) v0.1 pipeline and the model-based analysis for ChIP-Seq (MACS) v1.4.1 peak finding algorithm^68^ with a p-value enrichment threshold of 10^−9^ to define enhancers and super enhancers. We then used the BEDtools suite^69^ to compare the coincidence of enhancers and super enhancers with our unfused hypPB calling cards insertion peaks (**Fig 2** **and Fig 4SC-D**).

Additionally, for qualitative measures of histone modification enrichment at calling cards BRD4 insertion peaks (**Fig 2**), we used publicly available P0 mouse forebrain ChIP-Seq datasets from ENCODE^69^; specifically, H3K27ac (ENCSR094TTT), H3K4me1 (ENCSR465PLB), and H3K27me3 (ENCSR070MOK).

### Super enhancer in vitro sensitivity and specificity

Sensitivity and specificity of calling cards peaks were assessed for super enhancer identification in N2a cells that were either transfected or electroporated with AAV∷hypPB and AAV∷SRT plasmids (**Fig 4S**). RNA was separately isolated from a total of 33 wells (i.e. barcodes) from 6-well plates, and SRTs were sequenced, generating 806,653 unique insertions, though of note, the majority (651,631) were derived from 12 barcodes that received plasmids via electroporation. 800,000 unique insertions were randomly selected from the total pool of 806,653 insertions, from which significantly enriched peaks were defined using our background-free peak calling method at a range of significance thresholds. These peaks were intersected (with BEDtools intersect) with known N2a super enhancers defined via a previously published N2a H3K27ac ChIP-seq dataset^12^, and sensitivity was defined as the percentage of peaks intersecting super enhancers for each peak calling significance threshold. To then define specificity, we identified the “true negative” space of the genome, and assessed the percentage of true negative peaks intersected by calling cards peaks. To do this, we first identified any possible active enhancer region of the genome with MACS peak finding using a low-stringency significance threshold of 10^-1^ and subtracted these peaks from the mouse genome, creating a “true negative” genome. We then sampled peaks (with BEDtools shuffle) within this true negative genome of the same size distributions as the list of active super enhancers until we collected an average of 1X coverage across the genome. With a true negative space of 2,616,503,093 basepairs and a total super enhancer size of 23,143,876 basepairs, this required 114 random samplings, resulting in 85,158 true negative peaks. Finally, we intersected our calling card peaks with these true negative peaks, and specificity was defined as the percentage of true negative peaks not intersected by a calling card peak. Of note, we expect that unfused hypPB is driven to super enhancers via interaction with BRD4; thus, sensitivity and specificity measurements may be higher if compared to BRD4 occupancy rather than H3K27ac.

### Analysis of enhancer- and promoter-associated gene expression

Gene expression has been shown to be preferentially regulated by proximal enhancer elements^4,67,70^. Thus, since a cell type-specific mapping of enhancers to the genes they regulate is not available, we used proximity as an imperfect^6^ albeit widely used^3^ proxy. In our analyses of cell type-specific expression of genes near cell type-enriched BRD4 calling cards peaks (**Fig 3, Fig5S**), we first defined the nearest gene (or genes, if multiple intersected a calling cards peak) to each significant calling cards peak. These gene sets were then filtered and the remaining genes were used for subsequent analyses. Gene sets were filtered as follows: 1) genes greater than 10,000 bases away from a differential insertion peak were removed, to eliminate low confidence gene-enhancer pairs, 2) genes near or overlapping multiple insertion peaks counted once, and 3) genes for which cell type-specific RNA expression data were unavailable in our comparison dataset were removed.

Unbiased cell type identification was completed with the Cell-type Specific Expression Analysis (CSEA) tool^44^ (http://genetics.wustl.edu/jdlab/csea-tool-2/) using candidate gene sets near either GFAP∷Cre enriched or Syn1∷Cre enriched insertion peaks. For each set, we analyzed genes near the most enriched peaks for each cell type. For GFAP∷Cre, this included 131 genes (p<10^−21^ for associated insertion peaks), of which 114 were present in CSEA reference sets and used for analysis. For Syn1:Cre, this included 123 genes (p<10^−11^), with 110 present in reference sets.

For comparison of SP1 binding and gene expression in **Fig 5A-B**, we utilized the mm10_knownCanonical gene set and mm10_TSS coordinates from the UCSC genes table. We defined promoter-proximal regions as +/−1000 bases from the TSS. We first filtered mm10_knownCanonical gene set to remove duplicates (<3% of total genes) and then intersected gene coordinates with promoter proximal regions. After manually filtering to assign true promoters to each transcript (i.e. immediately upstream from TSS), we generated a list of unique promoter/gene combinations (24,528 unique genes) and compared insertion density and gene expression at these coordinates.

For comparison of P28 and P10 SP1 promoter insertions to RNA expression in **Fig 5C-G**, we utilized a previously published RNA-seq dataset^53^ with RNA expression data available for week 1 (Wk1) and week 4 (Wk4), which correspond to ~P7 and ~P28, respectively. Before assessing P28 or P10 insertion density at promoters, insertion profiles were downsampled such that each cohort had exactly 240,000 insertions per library (80,000 per mouse for P10, 3 mice; 60,000 per mouse for P28, 4 mice); thus, insertion totals could be directly compared without any normalization to library size. Further, this downsampling procedure eliminates the possibility that any given observed increase in insertion density at P28 was due to an overall increase in insertion total over time. We then calculated number of insertions at each unique promoter (using the list of unique promoter/gene combinations generated above) and removed any gene with no insertions at either timepoint (19,046 / 24,528 unique genes remaining). A pseudocount of 1 was added to promoter insertion totals for each gene at each timepoint prior to analysis. To eliminate noise due to low RNA expression and/or random low-frequency insertion events, we next removed any gene with <6 insertions combined between the P28 and P10 datasets (including the 2 pseudocounts) or <1 FPKM combined between Wk1 and Wk4 RNA-seq expression, leaving a final total of 4991 unique gene/promoter combinations which were used in subsequent analyses. This list of 4991 genes were divided into three categories, based on their RNA expression at Wk1 and Wk4: (1) early genes; log(Wk4/Wk1 FPKM) < −0.5, (2) constitutive genes; −0.5 < log(Wk4/Wk1 FPKM) < 0.5, and (3) late genes; log(Wk4/Wk1 FPKM > 0.5. Within these categories, SP1 occupancy was compared between the P28 and P10 cohorts.

### Validation of astrocyte enhancer candidates with PALE^46^

Candidate astrocyte-enriched enhancers were selected from the list of GFAP∷Cre-enriched insertion peaks in **Fig 3** based on three criteria, which select for high-confidence astrocyte enhancers: 1) enhancer size (smaller than 3kB, for cloning purposes), 2) significant GFAP∷Cre enrichment (over Syn1∷Cre), and 3) astrocyte-specific RNA expression of their nearest genes. These candidates were PCR amplified with primers listed in the table below and with an MluI overhang adapter (TGTAGGACGCGT) on either end, and cloned into the miniP-dsRed plasmid with MluI, upstream of the hsp68 minimal promoter. As a positive control, the canonical GFAP promoter^45^ was also cloned into this plasmid, in the same location.

To test efficacy of the candidates for enhancing dsRed expression, each plasmid was electroporated into lateral ventricle-proximal cells, along with a separate plasmid containing CFP driven by the canonical GFAP promoter. P0-1 pups were placed on a wet towel on wet ice for 10 minutes to anesthetize. Then plasmids were delivered to the lateral ventricle via intraventricular injection along with Fast Green FCF dye (Sigma, 2353-45-9), with coordinates approximately equidistant from the lambdoid suture and the eye, 2mm lateral to the sagittal suture, and 2mm depth to ensure lateral ventricle penetration. 1μl of DNA was delivered into one hemisphere for each mouse, at a concentration of 1μg/μl for GFAP∷CFP and 0.5μg/μl for dsRed plasmids. Electroporation was induced for 5 square pulses, 50ms per pulse at 100V and 950ms inter-pulse intervals, sweeping the electrodes from the dorsal to lateral using ~25° angle intervals.

Pups were sacrificed at P7 or P21 and brains were collected and analyzed with immunohistochemistry for CFP and dsRed (CFP and dsRed stained with GFP and RFP antibodies, respectively, as indicated in methods). Individual astrocytes from 3 brains (34-42 cells) per condition were imaged for intensity analysis in **Fig 3E-F** and **Fig 6SF,H** using equivalent exposure settings. A region of interest (ROI) was defined around the astrocyte based on CFP only, and dsRed and CFP fluorescence was quantified within the ROI using ImageJ (v. 1.51s). The ratio of CFP:dsRed was calculated for each cell and averaged and compared across conditions for final assessment of dsRed enhancement. For P7 co-localization quantifications in **Fig 6SG**, dsRed(+) cells were manually counted from 3 brains per condition, while P21 quantifications in **Fig 7B** were taken from multiple sections from a single brain per condition.

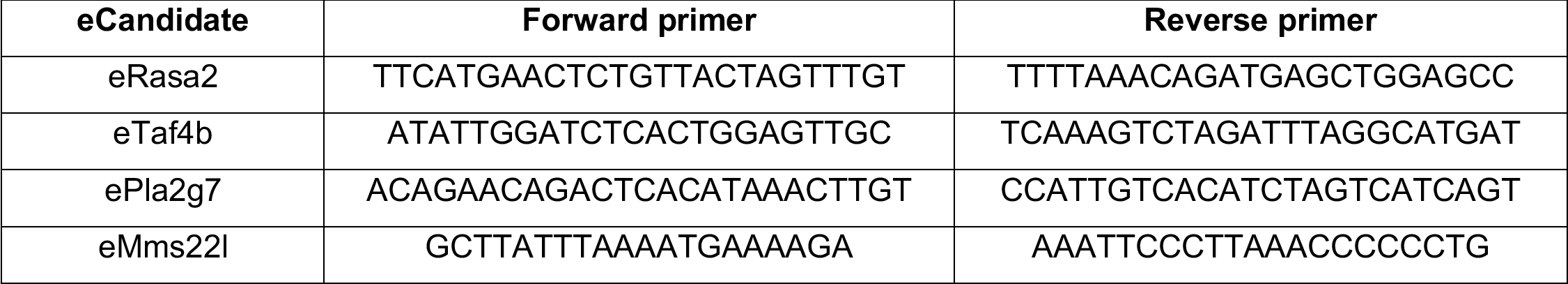

#### Statistical analyses

Statistical tests were done with GraphPad Prism v8.1.2 and are detailed in figure legends. For box-and-whisker plots, central tendency line represents median, box represents 25^th^-75^th^ percentiles, and whiskers represent minimum and maximum values.

#### Data availability

All raw and processed data is available through GEO accession GSE128493 (https://www.ncbi.nlm.nih.gov/geo/query/acc.cgi?acc=GSE128493) and the secure token for reviewer-only access has been provided to the editorial staff. Figures containing raw data include: 2, 3, 4, 5, 2S, 4S, 5S, and 6S. Supplemental tables containing information about significant peaks and figure-specific analyses are also available upon request.

#### Code availability

Calling card analysis software were developed previously^27^ and are available upon request.

