## Supplementary figures and legends for "A viral toolkit for recording transcription factor-DNA interactions in live mouse tissues"

**Supplemental figure legends**

*Figure 1S. AAV9 intracranial injection induces hypPB expression in the cortex.*

(A) *In situ* hybridization of hypPB RNA and no probe control displaying widespread cortical expression of hypPB after delivery of AAV::hypPB.

*Figure 2S. Comparison of neonatal and adult AAV delivery of traditional DNA calling card and SRT calling card systems.*

(A) Viral donor transposon constructs for DNA (AAV::BrokenHeart) and RNA (Self-Reporting Transposon; SRT) calling cards. (B) DNA calling cards library preparations were carried out as previously described in Wang et al., *Genetics*, 2012. TF-hypPB fusions insert BrokenHeart transposon DNA near TF binding sites. Genomic DNA is then harvested and digested with restriction enzymes that cut near the end of the transposon and in downstream genomic sequence. These fragments are subsequently self-ligated and circularized. From these, transposons and their flanking genomic sequences are amplified with inverse PCR, using primers that contain Illumina sequencing primers and adapters. Final products are sequenced and aligned to the mouse genome to map transposons to genomic locations. (C) Schematic of RNA calling cards library preparation protocol. HypPB inserts SRTs near TF binding sites, which then transcribe and report their genomic locations via RNA. RNA is reverse transcribed and PCR amplified, and Illumina adapters are added for sequencing. (D) Read mapping rates for BrokenHeart DNA calling cards and SRT RNA calling cards libraries after P0-1 or adult cortical delivery, showing increased efficiency of recovery for genome-mapping reads in SRT relative to BrokenHeart. (n=3 mice for P0-1 BrokenHeart [12.3% mapping to mm10]; n=2 mice for P0-1 SRT [80.7%]; n=3 mice for adult SRT [76.5%]) (E) Number of unique insertions observed from BrokenHeart DNA calling cards and SRT RNA calling cards libraries (n=3 mice for P0-1 BrokenHeart [198,981 insertions were recovered at mean read coverage of 8.98 reads/insertion]; n=2 mice for P0-1 SRT [3,732,694 insertions at 3.84 reads/insertion]; n=3 mice for adult SRT [4,114,106 insertions at 5.92 reads/insertion]), displaying greatly increased recovery with SRT. (F) Mapping locations of transposon insertions in various genomic regions from BrokenHeart and SRT calling card libraries.

*Figure 3S. AAV SRT calling cards system does not induce excess degeneration or behavioral/developmental deficits.*

(A-I) Behavioral and developmental assessments of mice injected at P0-1 with SRT calling cards (n=21) or control, RFP only (n=24) viruses revealed few or no developmental, sensorimotor, or anxiety-related deficits in calling card animals relative to control. All group comparisons were done with two-tailed, unpaired Student's t-test, with Bonferroni corrected $\alpha$=0.05 as a significance threshold (including all tests in Fig 1). **p<0.01 (Bonferroni corrected).

*Figure 4S. In vitro and in vivo sensitivity of unfused hypPB calling card libraries for active REs.*
(A-B) Sensitivity and specificity of super enhancer (SE) identification for unfused hypPB calling cards libraries in N2a cells. (A) Receiver-operator characteristic (ROC) curve for identification of active super enhancers using unfused hypPB peaks called from 800,000 unique insertions. Area under ROC curve: 0.82. (B) Super enhancer sensitivity at various significance thresholds and insertion totals, demonstrating high super enhancer sensitivity at even very low ($10^4$) insertion totals. (C-D) Intersections between significantly-enriched insertion peaks derived from *in vivo*, cortical unfused hypPB calling cards libraries and H3K27ac-marked (C) enhancers or (D) super enhancers at a range of insertion peak significance thresholds. Red line represents total number of significant peaks at each p-value threshold. Quantifications in Fig. 2F-I represent intersections at $p=10^{-30}$ significance threshold.

*Figure 5S. Supplemental to: FLEX calling cards system generates cell type-specific RE profiles.*
(A) Transfection of BrokenHeart transposons, hypPB FLEX plasmid, and Cre recombinase into HEK293T cells. n=3 wells per condition, 5 image fields per well, representative images shown. No TdTomato reporter from reconstituted BrokenHeart transposons observed in the absence of Cre or hypPB FLEX. (B) Representative images of Syn1::Cre and GFAP::Cre positive and negative littermate brains, displaying increased SRT-derived TdTomato reporter signal in Syn1::Cre positive animals. (C) Representative images displaying preferential expression in Neu(+) neurons in CA3 hippocampal regions of Syn1:Cre(+) mice, but not negative littermates. TdTomato images taken at equal exposure times for direct comparison. (D) Quantifications of unique insertions in Syn1::Cre or GFAP::Cre positive and negative littermate mice at sequencing depth downsampled to equal read depth and normalized to AAV::SRT viral titer (3X average coverage per insertion; two-tailed, unpaired Student's t-test: * $p<0.05$, **** $p<0.0001$). Syn1::Cre (+) vs (-): $p<0.0001$, $t=23.33$, $df=4$, 95% $C.I._{diff}$ [628,999 : 798,948]; GFAP::Cre (+) vs (-): $p=0.013$, $t=3.02$, $df=10$, 95% $C.I._{diff}$ [54,866 : 364,052] (E) Number of genes near differentially enriched insertion peaks in Syn1::Cre and GFAP::Cre animals at a range of significance thresholds. (F) Normalized neuron-to-astrocyte expression ratio [Neuron FKPM/(Neuron FPKM + Astrocyte FKPM] for genes near Syn1::Cre or GFAP::Cre enriched peaks at a range of significance thresholds for defining differentially enriched insertion peaks. As significance threshold becomes more stringent, expression of nearby genes becomes more cell type-specific. RNA-seq expression from Zhang et al., *Journal of Neuroscience*, 2014. (G) Astrocyte versus neuron expression of all genes near Syn1::Cre or GFAP::Cre enriched insertion peaks at a stringent peak-calling significance threshold ($p=10^{-7}$). Percent of genes on either side of the y=x midline shown. (H-I) Graphical representation of cortical cell type enrichment based on gene sets near either (H) Syn1::Cre ($p<10^{-11}$; top 123 genes) or (I) GFAP::Cre ($p<10^{-21}$; top 131 genes) enriched insertion peaks. Legend displays Benjamini-Hochberg corrected Fisher's Exact Test p-value for overlap of reference cell type-specific gene sets and Syn1::Cre or GFAP::Cre candidate gene sets. Stringencies for enrichment for each pre-defined reference set are represented by size of each hexagon, with the outer ring being the least stringent set and inner ring being the most stringent set.

*Figure 6S. Functional validation of enhancer activity at P7 for GFAP::Cre enriched unfused hypPB peaks.*

(A-D) Candidate GFAP::Cre enriched insertion peaks that were chosen for functional enhancer validation based on enhancer size, significant GFAP::Cre enrichment, and astrocyte-specific RNA expression of their nearest genes. Each candidate RE (highlighted in blue) was separately cloned into a plasmid upstream of the hsp68 minimal promoter and a dsRed reporter gene for *in vivo* testing. (E) Chromosomal coordinates and lengths of candidate REs. (F) Candidate RE reporter constructs were co-delivered along with a GFAP::CFP plasmid to ventricle-proximal radial glia, including astrocytes, via PALE[46]. Expression of dsRed was enhanced by both the canonical GFAP promoter (pGFAP; positive control) and three of the four candidate REs (all but eMms22l). (G) Percentage of dsRed(+) cells in cortex co-labeled with GFAP::CFP, demonstrating that >96% of labeled cells are glia at this timepoint in all conditions. n=150 dsRed(+) cells from 3 brains per condition. (H) Quantification of dsRed expression enhancement in GFAP::CFP(+) astrocytes by pGFAP and candidate RE constructs. n=34-42 GFAP::CFP(+) cells from 3 brains per condition (one-way ANOVA with Dunnett's multiple comparisons test; *p<0.05; **p<0.01; ****p<0.0001). $pGFAP_{mean\ diff.}$ = 0.66, 95% $C.I._{diff}$ [0.01 : 1.33]. $eRasa2_{mean\ diff.}$ = 0.67, 95% $C.I._{diff}$ [0.03 : 1.32]. $eTaf4b_{mean\ diff.}$ = 0.84, 95% $C.I._{diff}$ [0.21 : 1.48]. $ePla2g7_{mean\ diff.}$ = 1.22, 95% $C.I._{diff}$ [0.58 : 1.86]. $eMms22l_{mean\ diff.}$ = 0.002, 95% $C.I._{diff}$ [-0.65 : 0.65].

*Figure 7S. Candidate astrocyte enhancers identified with FLEX calling cards direct cell type-specific expression in astrocytes by P21.*

(A-B) Candidate astrocyte enhancers derived from FLEX calling cards enrichment (see Fig 6S) were electroporated into P0-1 mouse pups via PALE[46] and animals were sacrificed at P21 for IF analysis. In animals receiving a plasmid containing dsRed driven by a minimal promoter only (hsp68::dsRed), dsRed expression in the cortex was evident in a population of NeuN(+) neurons (white arrows) that was not observed at P7 (see Fig 6SF-G). In contrast, in animals receiving either the pGFAP-driven positive control plasmid or plasmids containing the candidate enhancers, dsRed expression was significantly limited to GFAP::CFP(+) cells, indicating that the enhancer candidates facilitate cell type-specificity of gene expression. n=67-201 dsRed(+) cells from a single brain per condition (one-way ANOVA with Dunnett's multiple comparisons test; ****p<0.0001). $pGFAP_{mean\ diff.}$ = 74.47%, 95% $C.I._{diff}$ [69.66% : 79.28%]. $eRasa2_{mean\ diff.}$ = 73.57%, 95% $C.I._{diff}$ [68.76% : 78.38%]. $eTaf4b_{mean\ diff.}$ = 74.47%, 95% $C.I._{diff}$ [69.66% : 79.28%]. $ePla2g7_{mean\ diff.}$ = 72.37%, 95% $C.I._{diff}$ [67.56% : 77.18%]. $eMms22l_{mean\ diff.}$ = 74.47%, 95% $C.I._{diff}$ [69.66% : 79.28%].

Figure 1S

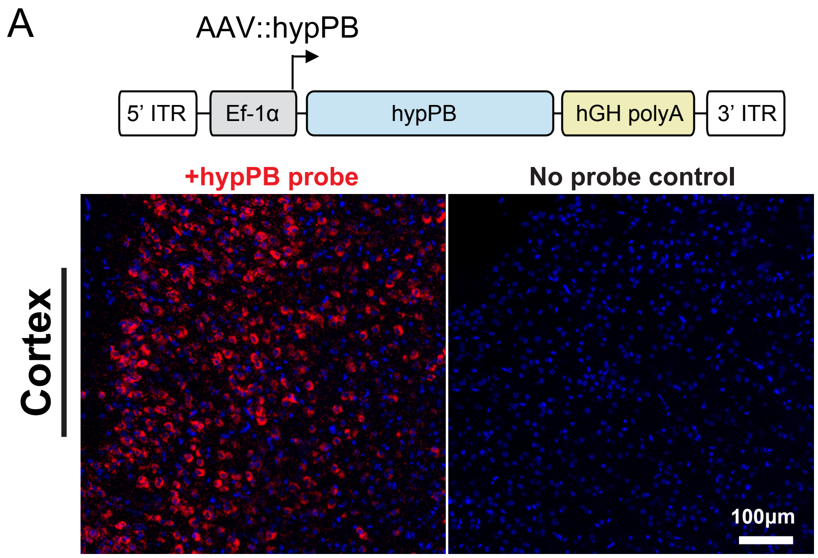
A
AAV::hypPB
5' ITR
Ef-1α
hypPB
hGH polyA
3' ITR
+hypPB probe
No probe control
Cortex
100μm

Figure 2S

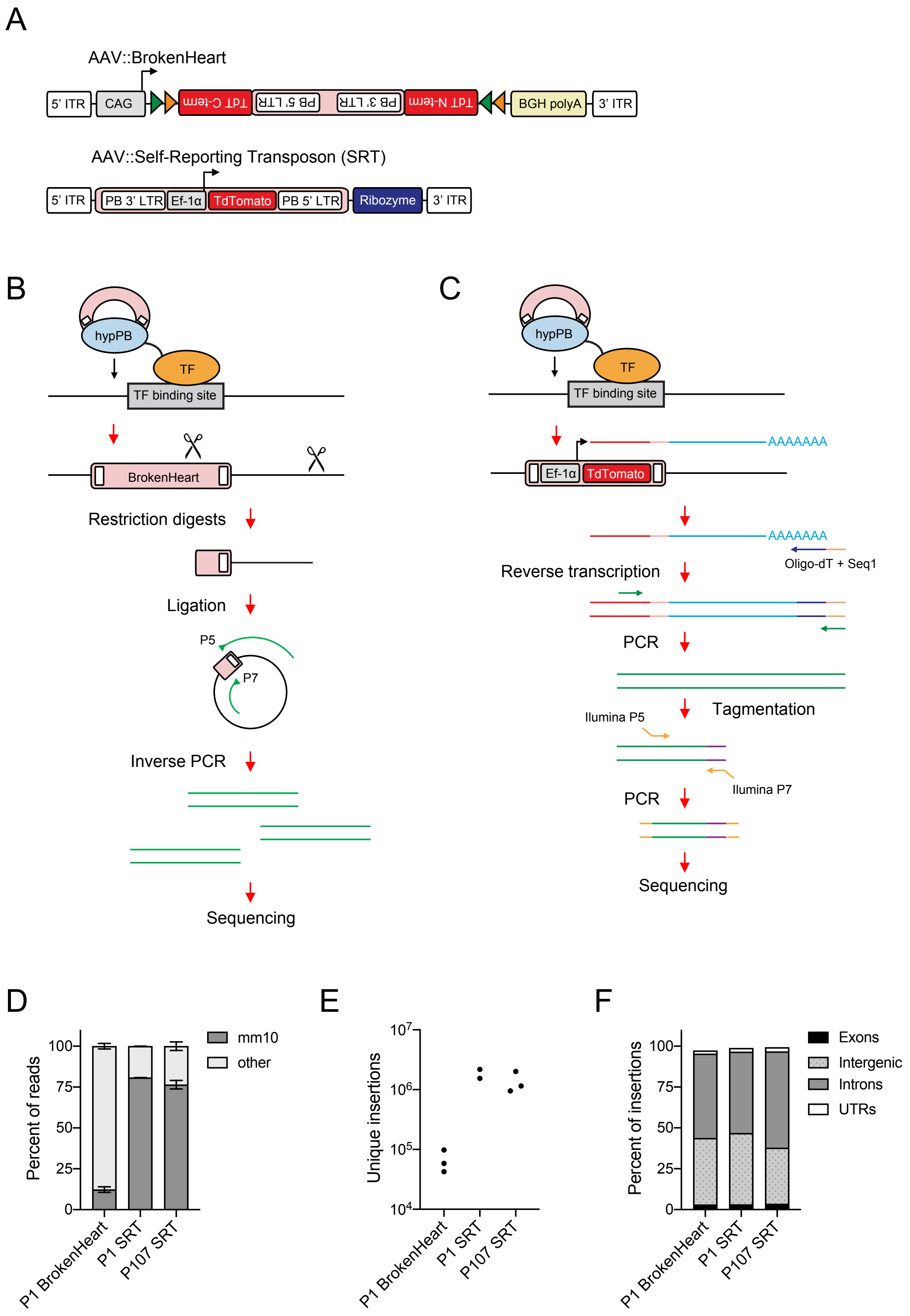
A
AAV::BrokenHeart
5' ITR
CAG
TdT C-term
PB 5' LTR
PB 3' LTR
TdT N-term
BGH polyA
3' ITR
AAV::Self-Reporting Transposon (SRT)
5' ITR
PB 3' LTR
Ef-1α
TdTomato
PB 5' LTR
Ribozyme
3' ITR
B
hypPB
TF
TF binding site
BrokenHeart
Restriction digests
Ligation
P5
P7
Inverse PCR
Sequencing
C
hypPB
TF
TF binding site
Ef-1α
TdTomato
AAAAAAA
AAAAAAA
Oligo-dT + Seq1
Reverse transcription
PCR
Tagmentation
Ilumina P5
Ilumina P7
PCR
Sequencing
D
Percent of reads
100
75
50
25
0
mm10
other
P1 BrokenHeart
P1 SRT
P107 SRT
E
Unique insertions
$10^7$
$10^6$
$10^5$
$10^4$
P1 BrokenHeart
P1 SRT
P107 SRT
F
Percent of insertions
100
75
50
25
0
Exons
Intergenic
Introns
UTRs
P1 BrokenHeart
P1 SRT
P107 SRT

### Figure 3S

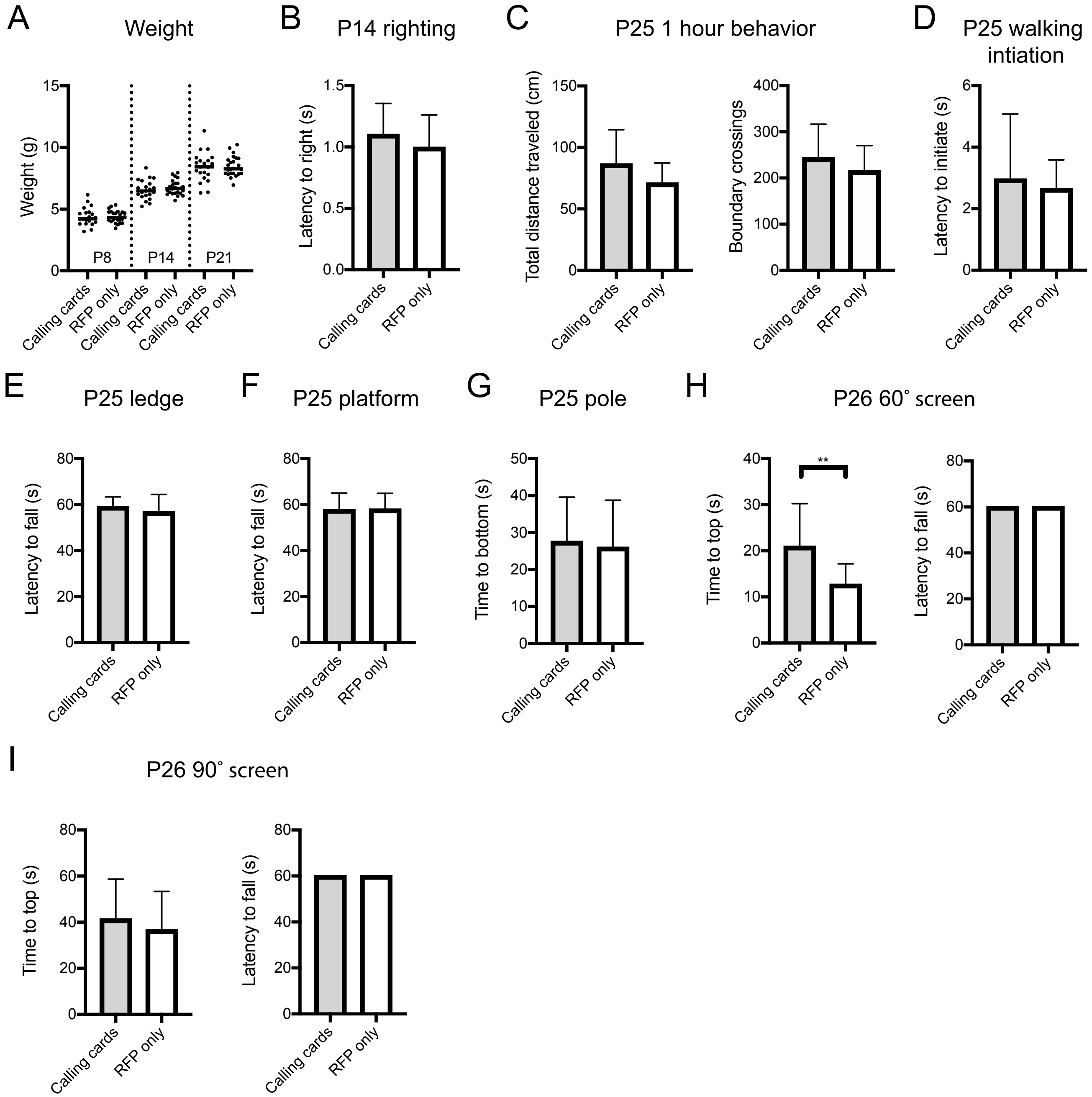
A
Weight
Weight (g)
P8
P14
P21
B
P14 righting
Latency to right (s)
C
P25 1 hour behavior
Total distance traveled (cm)
Boundary crossings
D
P25 walking intiation
Latency to initiate (s)
E
P25 ledge
Latency to fall (s)
F
P25 platform
Latency to fall (s)
G
P25 pole
Time to bottom (s)
H
P26 60˚ screen
Time to top (s)
**
Latency to fall (s)
I
P26 90˚ screen
Time to top (s)
Latency to fall (s)
Calling cards
RFP only

Figure 4S

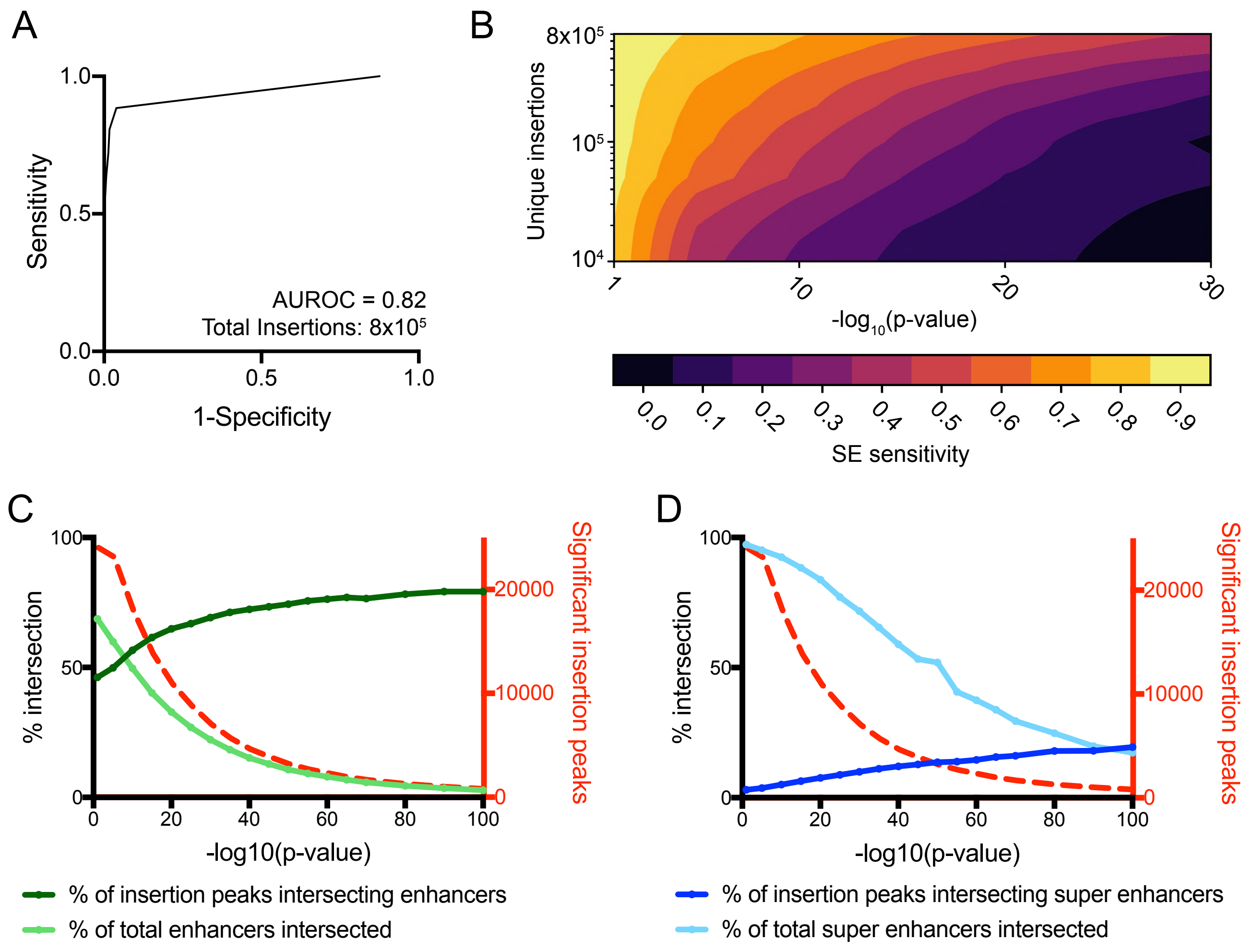
A
1.0
0.5
0.0
Sensitivity
0.0
0.5
1.0
1-Specificity
AUROC = 0.82
Total Insertions: 8x10⁵
B
8x10⁵
10⁵
10⁴
Unique insertions
1
10
20
30
-log₁₀(p-value)
0.0
0.1
0.2
0.3
0.4
0.5
0.6
0.7
0.8
0.9
SE sensitivity
C
100
50
0
% intersection
20000
10000
0
Significant insertion peaks
0
20
40
60
80
100
-log10(p-value)
% of insertion peaks intersecting enhancers
% of total enhancers intersected
D
100
50
0
% intersection
20000
10000
0
Significant insertion peaks
0
20
40
60
80
100
-log10(p-value)
% of insertion peaks intersecting super enhancers
% of total super enhancers intersected

### Figure 5S

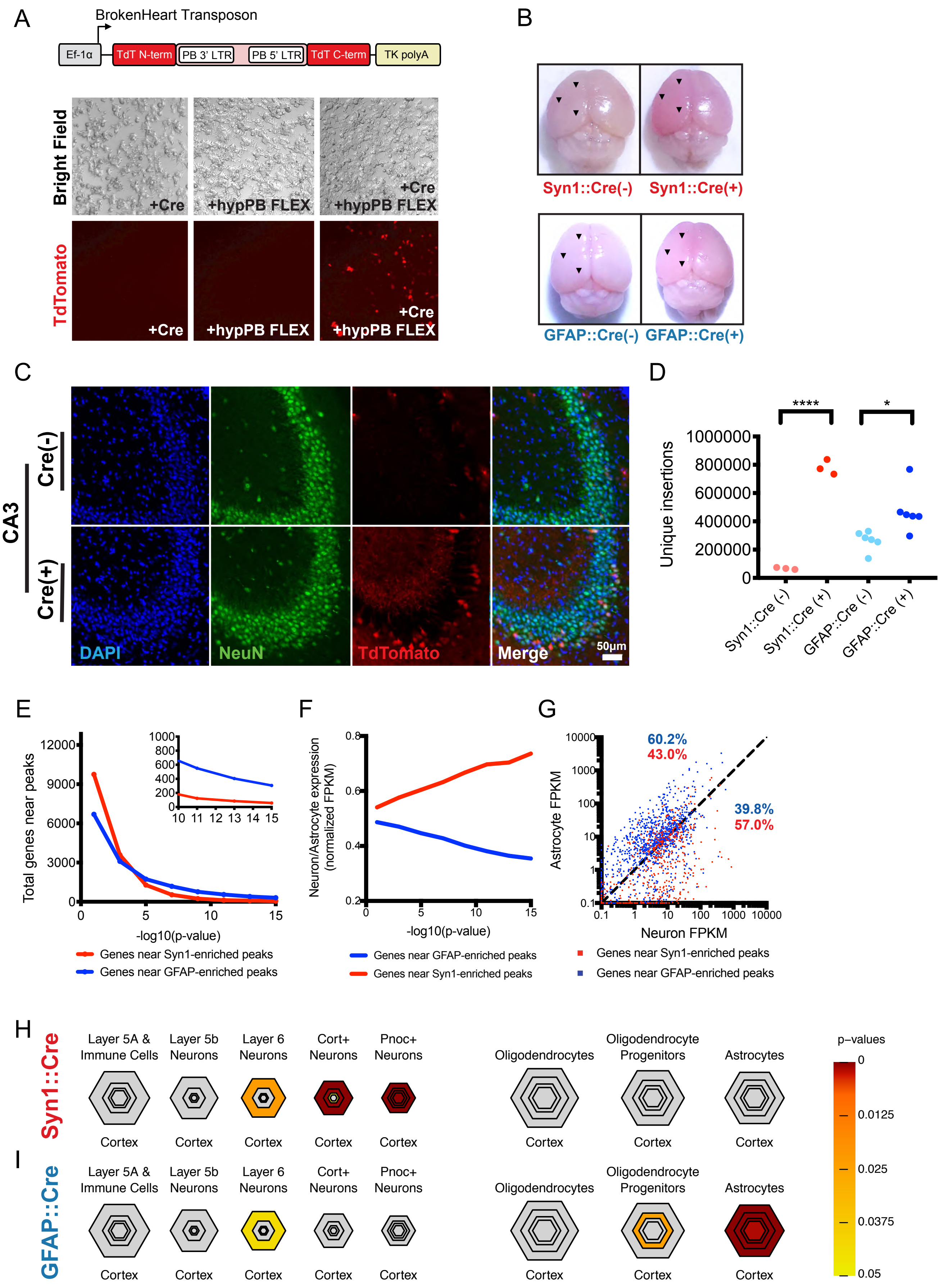
A
BrokenHeart Transposon
Ef-1α
TdT N-term
PB 3' LTR
PB 5' LTR
TdT C-term
TK polyA
Bright Field
TdTomato
+Cre
+hypPB FLEX
+Cre +hypPB FLEX
B
Syn1::Cre(-)
Syn1::Cre(+)
GFAP::Cre(-)
GFAP::Cre(+)
C
CA3
Cre(-)
Cre(+)
DAPI
NeuN
TdTomato
Merge
50μm
D
Unique insertions
****
*
Syn1::Cre (-)
Syn1::Cre (+)
GFAP::Cre (-)
GFAP::Cre (+)
E
Total genes near peaks
-log10(p-value)
Genes near Syn1-enriched peaks
Genes near GFAP-enriched peaks
F
Neuron/Astrocyte expression (normalized FPKM)
-log10(p-value)
Genes near GFAP-enriched peaks
Genes near Syn1-enriched peaks
G
Astrocyte FPKM
Neuron FPKM
60.2%
43.0%
39.8%
57.0%
Genes near Syn1-enriched peaks
Genes near GFAP-enriched peaks
H
Syn1::Cre
I
GFAP::Cre
Layer 5A & Immune Cells
Layer 5b Neurons
Layer 6 Neurons
Cort+ Neurons
Pnoc+ Neurons
Oligodendrocytes
Oligodendrocyte Progenitors
Astrocytes
Cortex
p-values
0
0.0125
0.025
0.0375
0.05

### Figure 6S

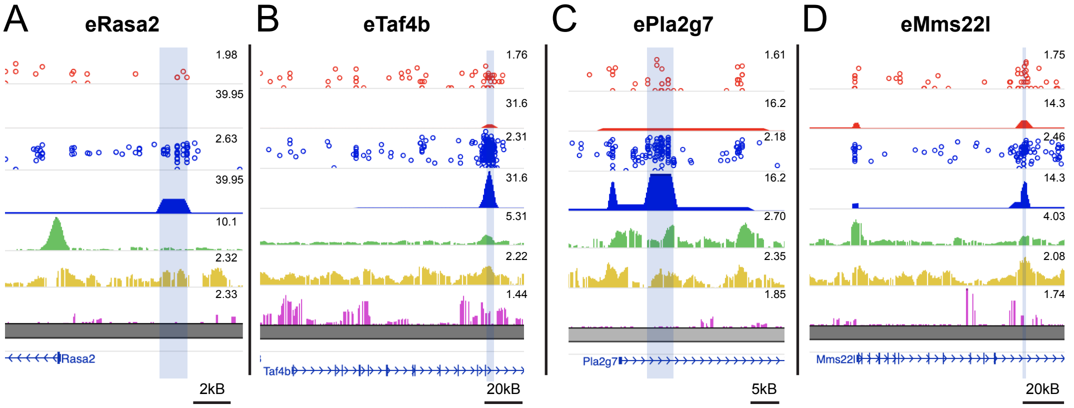


E

| Enhancer region | mm10 chrom. coordinates | Length (bp) |
|---|---|---|
| eRasa2 | chr9:96636934 -96638463 | 1530 |
| eTaf4b | chr18:14845190 -14846791 | 1602 |
| ePla2g7 | chr17:43571375 -43574084 | 2710 |
| eMms22l | chr4:24544409 -24544521 | 112 |

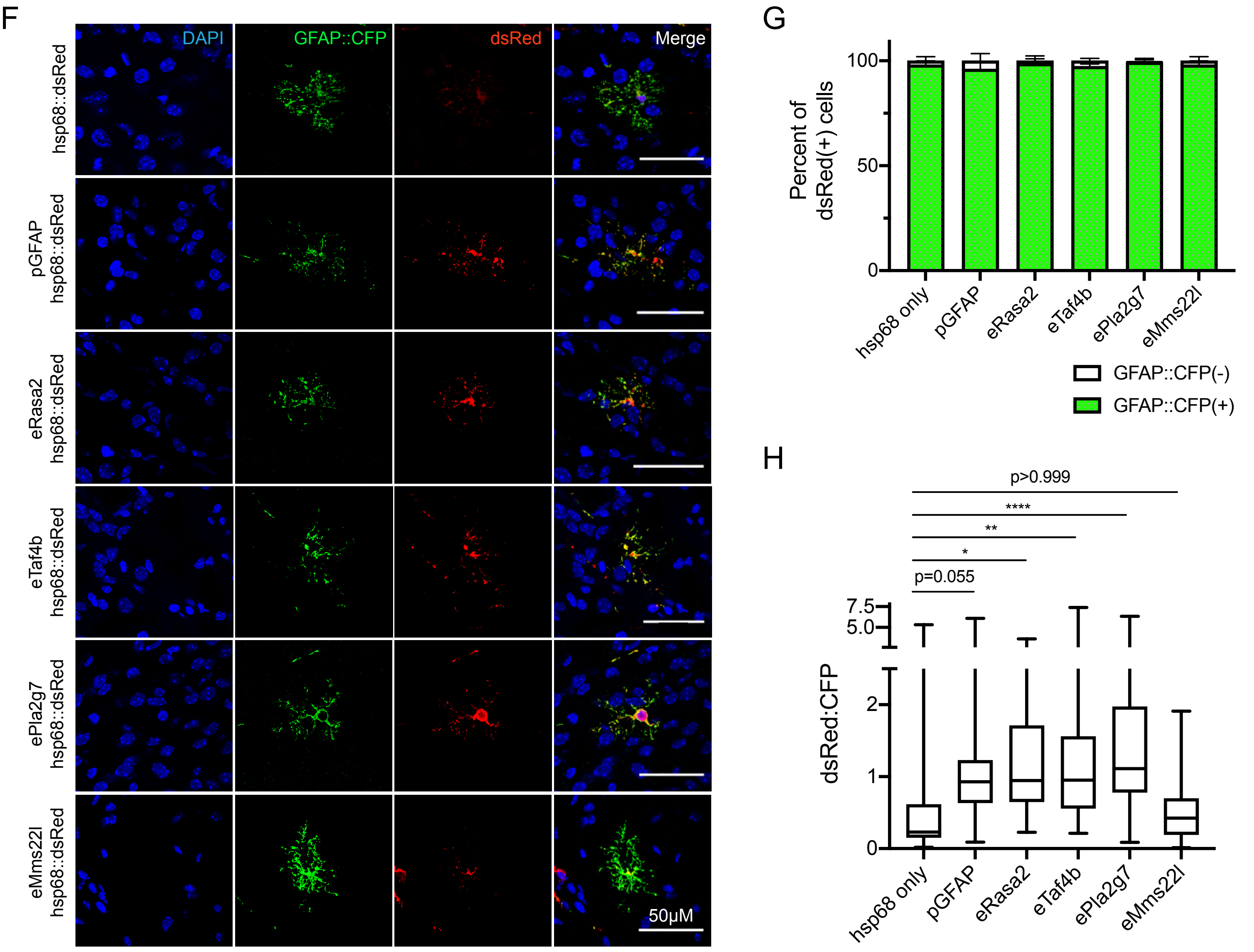

### Figure 7S

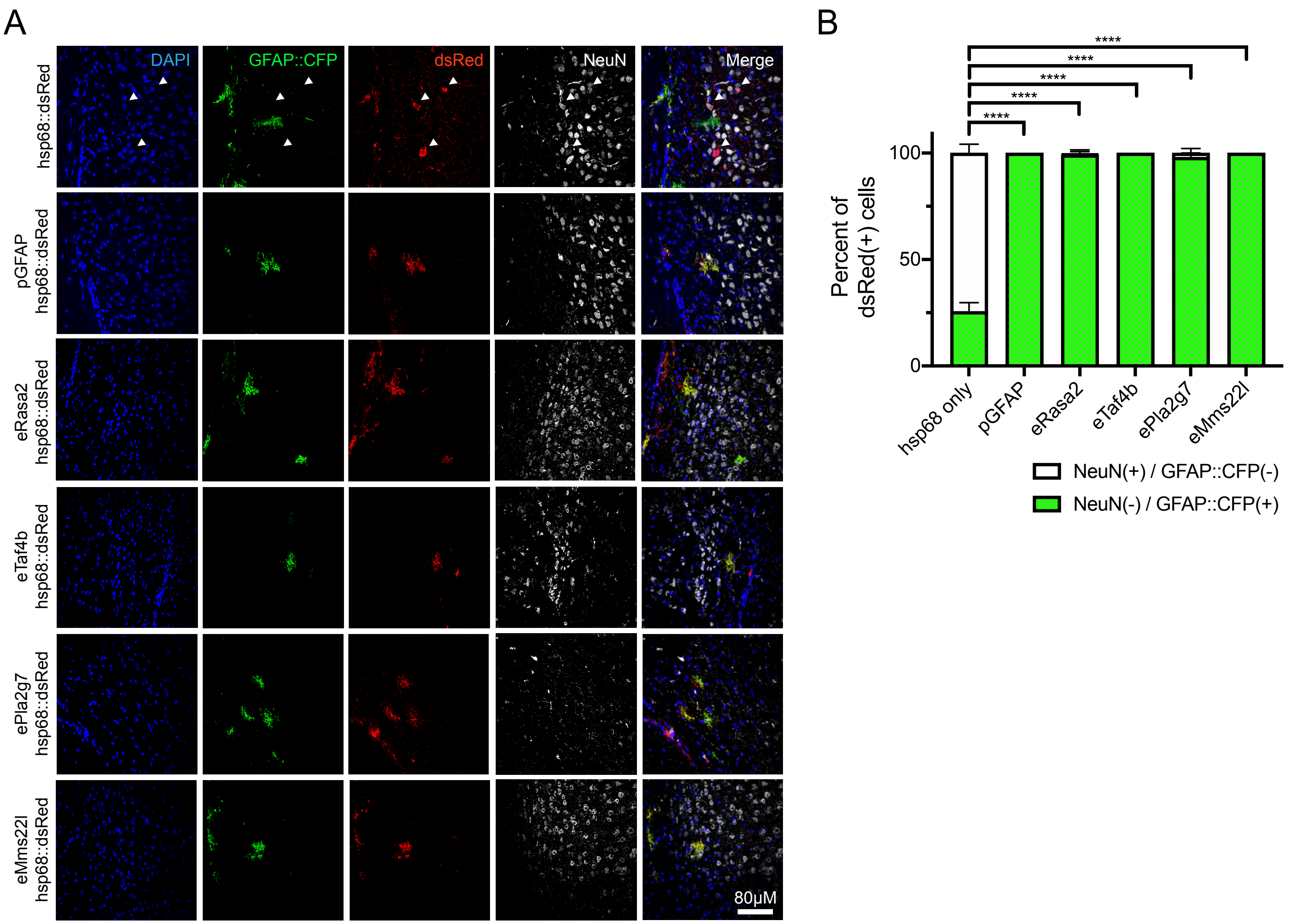
A
DAPI
GFAP::CFP
dsRed
NeuN
Merge
hsp68::dsRed
pGFAP hsp68::dsRed
eRasa2 hsp68::dsRed
eTaf4b hsp68::dsRed
ePla2g7 hsp68::dsRed
eMms22l hsp68::dsRed
80μM
B
Percent of dsRed(+) cells
0
50
100
****
****
****
****
****
hsp68 only
pGFAP
eRasa2
eTaf4b
ePla2g7
eMms22l
NeuN(+) / GFAP::CFP(-)
NeuN(-) / GFAP::CFP(+)
